# Adenylnucleotide-mediated binding of the PII-like protein SbtB contributes to controlling activity of the cyanobacterial bicarbonate transporter SbtA

**DOI:** 10.1101/2021.02.14.431189

**Authors:** Britta Förster, Bratati Mukherjee, Loraine M. Rourke, Joe A. Kaczmarski, Colin J. Jackson, G. Dean Price

## Abstract

Cyanobacteria have evolved a remarkably powerful CO_2_ concentrating mechanism (CCM), enabling high photosynthetic rates in environments with limited inorganic carbon (Ci). Therefore, this CCM is a promising system for integration into higher plant chloroplasts to boost photosynthetic efficiency and yield. The CCM depends on active Ci uptake, facilitated by bicarbonate transporters and CO_2_ pumps, to elevate CO_2_ concentration around the active sites of the primary CO_2_ fixing enzyme, Rubisco, which is encapsulated in cytoplasmic micro-compartments (carboxysomes). The essential CCM proteins have been identified, but the molecular signals and regulators that coordinate function in response to light, Ci availability and other environmental cues are largely unknown. Here, we provide evidence, based on a novel *in vitro* binding system, for a role of the PII-like SbtB protein in regulating Ci uptake by the bicarbonate transporter, SbtA, in response to the cellular adenylate energy charge (AEC) through dynamic protein-protein interaction. Binding of the SbtA and SbtB proteins from two phylogenetically distant species, *Cyanobium sp*. PCC7001 and *Synechococcus elongatus* PCC7942, was inhibited by high ATP, and promoted by low [ATP]:[ADP or AMP] ratios *in vitro*, consistent with a sensory response to the AEC mediated through adenylnucleotide ligand-specific conformation changes in SbtB. *In vivo*, cell cultures of S. elongatus showed up to 70% SbtB-dependent down-regulation of SbtA bicarbonate uptake activity specifically in the light activation phase during transitions from dark to low light when low cellular AEC is expected to limit metabolic activity. This suggests SbtB may function as a curfew protein during prolonged low cellular AEC and photosynthetically unfavourable conditions to prevent energetically futile and physiologically disadvantageous activation of SbtA.

## Introduction

The cyanobacterial CO_2_ concentrating mechanism (CCM) is one of the most efficient CCMs known to date. It has been fundamental to their adaptability and successful competition in environments providing limited inorganic carbon (Ci) supply for oxygenic photosynthesis and is reflected in an estimated contribution of 30-80% global primary productivity in the oceans (Liu et al., 1997; Field et al., 1998). Therefore, recent efforts to enhance photosynthetic carbon fixation in crop species have focussed on introducing a synthetic cyanobacterial CCM into chloroplasts, which has been predicted to translate into 36-60% higher crop productivity (Price et al., 2013; McGrath and Long, 2014; Long et al., 2016; Meyer et al., 2016). The cyanobacterial CCM enables active uptake of two Ci species, CO_2_ and bicarbonate (HCO_3_^-^), which leads to accumulation of a cytoplasmic bicarbonate pool as large as 20-40 mM (Badger and Andrews, 1982; Price and Badger, 1989). This bicarbonate pool supports carbonic anhydrase (CA)-mediated elevation of the CO_2_ concentration around the primary CO_2_-fixing enzyme ribulose-1,5-bisphoshate carboxylase/oxygenase (Rubisco) inside proteinaceous micro-compartments known as carboxysomes. Consequently, Rubisco carboxylation is enhanced over oxygenation, suppressing energetically costly photorespiration (for review see (Badger and Price, 1992; Kaplan and Reinhold, 1999; Price et al., 2008; Espie and Kimber, 2011; Price, 2011)). To date, five Ci uptake systems have been verified, varying with respect to energization, substrate affinity, flux rates and induction by low Ci conditions (Price, 2011). Two light-dependent, thylakoid-located CO_2_ pumps or energy-coupled CAs, Ndh-1_3_ and Ndh-1_4_ (Maeda et al., 2002; Ohkawa et al., 2002), convert external and internally recaptured CO_2_ to HCO_3_^-^, energized by NADPH and reduced ferredoxin (Ogawa et al., 1985; Maeda et al., 2002; Price et al., 2008). Three types of plasma membrane-located bicarbonate transporters, the ATP-dependent BCT1 complex (Omata et al., 1999) and the HCO_3_^-^/Na^+^ symporters BicA (Price et al., 2004) and SbtA (Shibata et al., 2002), transfer bicarbonate directly from the external environment into the cytoplasm upon activation by low Ci conditions and light (Sültemeyer et al., 1998; McGinn et al., 2003). Of particular interest for a synthetic CCM is SbtA due to its ability to function in nonphotosynthetic *E. coli* (Du et al., 2014). SbtA requires a Na^+^ gradient directed from the outside to the inside of the cell for activity (Shibata et al., 2002; Price et al., 2004). Both CO_2_ uptake and HCO_3_^-^ transport are activated by light within 5 and 30 s, respectively, (Badger and Andrews, 1982; Price et al., 2011). Early on, Kaplan *et al*. reported a gradual inactivation of Ci uptake in the dark, suggesting a link to the state of photosynthetic electron transport or a redox signal (Kaplan et al., 1987), but the molecular mechanisms regulating the activity of specific bicarbonate transporters are largely unknown.

A highly conserved feature of cyanobacterial genomes is the presence of the *sbtB* gene, encoding the PII-like SbtB protein, in the immediate gene neighbourhood of the *sbtA* gene (Rae et al., 2011). Both genes are co-expressed under Ci limitation in *Synechococcus elongatus* PCC7942 and *Synechocystis sp.* PCC 6803 (Woodger et al., 2003; Schwarz et al., 2011), suggesting a functional connection between the SbtA and SbtB proteins. This has been supported by evidence from co-expression of several SbtA and SbtB proteins in *E. coli*, where SbtA-mediated bicarbonate uptake was specifically inactivated in the presence of the cognate SbtB protein, and a physical interaction was shown for the SbtA and SbtB proteins from *S. elongatus* PCC7942 (hereafter SbtA_7942_ and SbtB_7942_, respectively) (Du et al., 2014). Similarly, a study by Selim *et al*. (Selim et al., 2018) indicated SbtA and SbtB from *Synechocystis sp.* PCC6803 (hereafter SbtA_6803_ and SbtB_6803_, respectively) interacted in response to Ci availability, which was enhanced by addition of ADP or AMP and diminished by cAMP. It was suggested the cAMP:AMP ratio may indirectly signal intracellular Ci levels and thus regulate SbtA_6803_:SbtB_6803_ complex formation, but effects on SbtA activity were not directly assessed. Meanwhile, a broader role for SbtB_6803_ in Ci and metabolic sensing has emerged. The same study (Selim et al., 2018) showed that the *sbtB6803* knockout mutant was impaired in acclimation to high Ci conditions, remaining locked in the high Ci affinity state typical of low Ci acclimation. Moreover, SbtB_6803_ was implicated in far red light induced inhibition of photosynthetic oxygen evolution (Oren et al., 2021) and in bis-(3′,5′)-cyclic diadenosine monophosphate (c-di-AMP) and redox state sensing linked to control of glycogen synthesis and growth performance of cyanobacterial cells during day/night cycles (Selim et al., 2021; Selim et al., 2023).

Structural evidence for SbtA:SbtB interaction comes from X-ray crystallography and cryo-electron microscopy, which resolved trimeric SbtA_6803_ bound to SbtB_6803_ trimers in the presence of AMP as a ligand to SbtB (Fang et al., 2021; Liu et al., 2021). Furthermore, SbtB proteins have been classified as non-canonical PII-like members of the multi-functional PII and PII-like protein superfamily that consists of transcriptional and protein activity regulators, environmental signal transducers or sensors, widely-distributed across bacteria, archaea and plants (Radchenko et al., 2010). In cyanobacteria, there is substantial evidence for a central role of PII proteins in sensing and balancing the carbon and nitrogen status of the cell as part of regulatory networks that affect Ci and N acquisition (Forchhammer and Selim, 2020). In crystals, SbtB proteins form homotrimers typical of PII with clefts between monomers forming binding sites for small effector molecules (Forchhammer and Lüddecke, 2016) that may bind in a competitive manner or cooperatively. Common ligands for PII proteins are ATP, ADP, Mg^2+^ and 2-oxoglutarate (2OG) which impact on protein conformation, particularly the T-loop, and the interaction with target proteins (Radchenko et al., 2010; Huergo et al., 2013; Zeth et al., 2014; Forchhammer and Lüddecke, 2016; Forchhammer and Selim, 2020). Ligands for cyanobacterial PII-like proteins include ADP and bicarbonate for the carboxysome-associated CPII from *Thiomonas intermedia* (Wheatley et al., 2016), and adenylnucleotides for the SbtB_6803_ protein, i.e. ATP, ADP, AMP, cAMP and c-di-AMP(Selim et al., 2018; Selim et al., 2021) and the SbtB protein from *Cyanobium sp.* PCC7001 (SbtB_7001_), i.e. cAMP, AMP, ADP, ATP and Ca^2+^ATP (Kaczmarski et al., 2019). SbtB_6803_ and SbtB_7001_ share the basic architecture, but differ in ligand-coordinating residues, ligand affinities and ligand-induced conformation changes. Recently, SbtB_6803_ was shown to exert apyrase activity, slowly converting ATP ligands to AMP in a redox state-dependent manner (Selim et al., 2023). Apyrase activity was tightly linked to the oxidation and disulfide bond formation between cysteines of the C-terminal R-loop, which is an extension that is present on a subset of SbtB proteins but not in SbtB_7001_ and SbtB_7942_. Therefore, one may expect fundamental mechanistic differences from SbtB_6803_.

The energy status of living cells is reflected in the relative abundance rather than absolute concentrations of the adenylnucleotides ATP, ADP and AMP and referred to as the cellular adenylate energy charge (AEC), described quantitatively as ([ATP]+0.5[ADP])/([ATP]+[ADP]+[AMP]) (Chapman et al., 1971). Intriguingly, SbtB_7001_ had 18 to 38-fold higher binding affinities for ATP over ADP, AMP and cAMP in the presence of Ca^2+^ (Kaczmarski et al., 2019), which suggested to us certain SbtB proteins may sense the AEC directly and dynamically interact with SbtA in response to changes in adenylate nucleotide ratios. We propose that the adenylnucleotide ligand-induced structural changes observed for SbtB proteins determine SbtA:SbtB complex formation and control of the SbtA activation state. Here, we demonstrate that the adenylnucleotides ATP, ADP, AMP and cAMP regulate *in vitro* binding of SbtA (SbtA_7001_) from *Cyanobium sp.* PCC7001 and SbtA_7942_ to their cognate SbtB proteins, and present *in vivo* effects of SbtB on SbtA activity in cyanobacteria linked to light-induced changes in AEC, consistent with a role of SbtB as AEC sensor and response regulator.

## Results

### SbtA:SbtB complex formation is affected specifically by adenylnucleotides

For SbtA:SbtB interaction studies, we co-expressed SbtA_7001_, or SbtA_7942_, with their cognate SbtB proteins which were C-terminally tagged with human influenza hemagglutinin (HA) epitope and 5 or 6 histidines (His) in the *E.coli* strain DH5α (*E. coli*) and in a bicarbonate transporter-deficient *Synechococcus elongatus* PCC7942 mutant (*Synechococcus* ΔCS). The SbtB_7001_ and SbtB_7942_ protein were identified immunologically at 13 kDa, consistent with their calculated molecular weights, but the apparent molecular weights of SbtA_7001_ and SbtA_7942_ monomers (23 kDa and 29 kDa, respectively) were 11 kDa lower than their calculated molecular weights, as is commonly observed for membrane proteins and attributed to altered protein-SDS detergent interactions (Rath et al., 2009). Specific binding of SbtA to SbtB proteins was evaluated *in vitro* by metal immobilized chromatography (IMAC) using membrane-enriched, native protein extracts that contained both SbtA and HAHis-tagged SbtB proteins (**Fig. S1**). Only the HAHis-SbtB but no SbtA protein was found to attach to Ni^2+^ coated beads which ensured co-elution of SbtA and SbtB from the beads was exclusively due to the interaction of the two proteins. The *in vitro* binding assays without addition of effectors showed that SbtA and SbtB were largely dissociated in native protein extracts (**Figs. 1, S2**), which led us to focus on the association of SbtA:SbtB complexes in response to a range of potential effector molecules and co-factors reported to interact with PII and PII-like proteins. Based on our previous observation that SbtB_7001_ binds ATP, ADP, AMP and cAMP with micromolar affinities (Kaczmarski et al., 2019), we expected SbtA:SbtB complex formation and dissociation be affected by these molecules. Since Ca^2+^ enhanced binding of ATP to SbtB_7001_ *in vitro* (Kaczmarski et al., 2019), we also included 200 nM Ca^2+^ in binding assays as a potential cofactor, which was equivalent to the basal Ca^2+^ levels in cyanobacterial cells (Leganés et al., 2009) did not facilitate SbtA:SbtB binding by itself (**Fig. S2**).

**Fig. 1.**
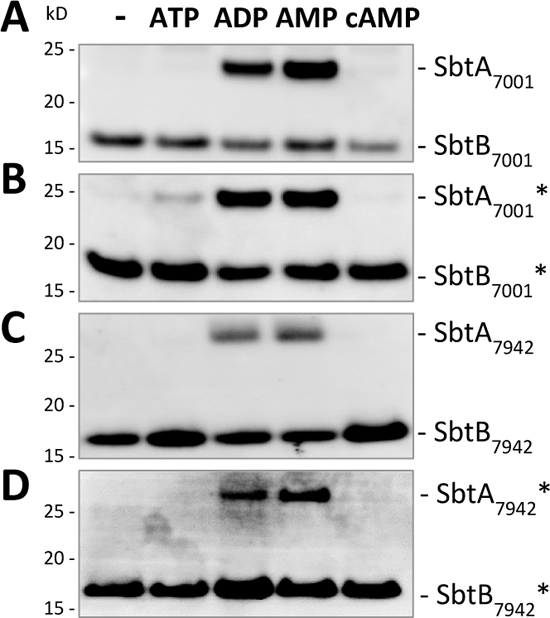
Association of SbtA and cognate SbtB proteins in response to adenylnucleotides. The SbtA_7001_-SbtB_7001_ and SbtA_7942_-SbtB_7942_ pairs were expressed in *E. coli* (**A, C**) and *Synechococcus* ΔCS (denoted by *; **B, D**). The HAHis-tagged SbtB proteins were immobilized to examine interaction with SbtA proteins and adenylnucleotides (2 mM ATP, ADP, AMP, or cAMP) during *in vitro* IMAC binding assays. The HAHis-SbtB protein and corresponding amounts of SbtA retained in SbtA:SbtB complexes were detected by immunoblot analyses. Representative results are shown of n=3-8 experiments per adenylnucleotide treatment.

Under these conditions, the association of stable SbtA_7001_:SbtB_7001_ and SbtA_7942_:SbtB_7942_ complexes was observed exclusively in the presence of 2 mM ADP or AMP (**Fig. 1**). Conversely, SbtA and SbtB remained fully dissociated in the presence of 2 mM ATP or cAMP (**Fig. 1**).

In addition, a potential link between stabilization of SbtA:SbtB complexes and photosynthetic activity was tested, but SbtA:SbtB binding was not affected by the Calvin-Benson-Basham (CBB) cycle intermediates ribulose-1,5-bisphosphate (RUBP), 3-phosphoglycerate (3PGA) (**Fig. 2**). Similarly, 2OG which is generated in the tricarboxylic acid (TCA) cycle and acts a master regulatory metabolite coordinating carbon and nitrogen metabolism in association with canonical PII proteins (Huergo and Dixon, 2015), did not facilitate SbtA:SbtB complex formation (**Fig. 2**).Combined with ATP or ADP, 2OG had no synergistic effect (**Fig. 2**), in contrast to the prerequisite binding of Mg^2+^ATP to the *E. coli* PII protein for 2OG to become effective (Radchenko et al., 2013). Furthermore, we investigated possible substrate-level regulation of SbtA activity via HCO_3_ ^-^-dependent complex formation with SbtB. HCO_3_ ^-^ had no effect on SbtA:SbtB binding irrespective of combinations with Ca^2+^ or Mg^2+^, ATP or AMP (**Fig. S2, S3**). In summary, we observed that small molecule effectors unable to bind to SbtB_7001_ (Kaczmarski et al., 2019), were equally ineffective in mediating the binding of either SbtA-SbtB pair. It is notable that SbtA-SbtB-effector interactions were very similar for proteins expressed in *E. coli* compared to *Synechococcus* ΔCS (**Fig. 1, 2**), which suggests proteins were not subject to species-dependent, post-translational modifications and validates use of *E. coli* as heterologous expression system.

**Fig. 2.**
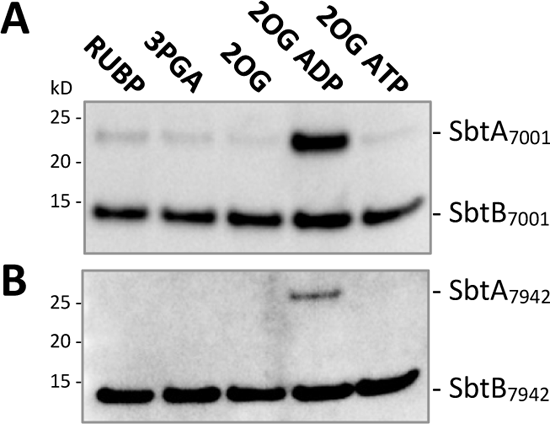
The association of SbtA and cognate SbtB proteins is not directly linked to key metabolites of the CBB cycle (RUBP, 3PGA) and the TCA cycle (2OG). *In vitro* IMAC binding assays were performed with the SbtA_7001_-SbtB_7001_ (**A**) and SbtA_7942_-SbtB_7942_ (**B**) pairs expressed in *E. coli* in the presence of 2 mM RUBP, 3PGA or 2OG, and 2 mM 2OG combined with 2 mM ADP, or ATP (2OG ADP and 2OG ATP, respectively). The representative immunoblot images show HAHis-tagged SbtB proteins and the SbtA proteins retained in SbtA:SbtB complexes; n=2.

### The AEC simulated by adenylnucleotide ratios determined the SbtA:SbtB complex formation

The fact that SbtA:SbtB complexes were destabilized by ATP and stabilized by either ADP or AMP, strongly suggested complex formation would be impacted by changes in the cellular adenylate energy charge (AEC). Therefore, we simulated defined energy charge environments *in vitro* with different molar ratios of ATP:ADP or ATP:AMP within a constant total adenylnucleotide pool of 2 mM. Both SbtA-SbtB pairs, irrespective of expression in *E. coli* or *Synechococcus* ΔCS, showed decreasing amounts of SbtA bound to SbtB with an increasingly higher proportion of ATP over ADP (**Fig. 3A-D**) and AMP (**Fig. 3E,F**). Since both SbtA-SbtB pairs were unbound in the presence of cAMP (**Fig. 1**), we replaced ATP with cAMP to test whether SbtA:SbtB binding could be affected by the molar ratios of cAMP to ADP or AMP, similar to what has been reported for the SbtA_6803_-SbtB_6803_ pair (Selim et al., 2018). It was indeed possible to impair SbtA:SbtB binding with high cAMP:ADP or AMP ratios (**Fig. 4**). However, to evaluate the effects of cAMP in relation to ATP, it is important to recognize that ATP concentrations up to 2 mM corresponded to the physiological range of intracellular ATP pools measured in several bacteria (Thauer et al., 1977; Yaginuma et al., 2014), whereas estimates of cAMP levels in cyanobacterial cells have been up to 1000-fold lower than the intracellular ATP levels, i.e. in the nano- to micromolar range (Ohmori and Okamoto, 2004). The *in vitro* cAMP levels reducing the amount of bound SbtA to 50% was 1.16 mM (**Fig. 5B**, AR_50_ of SbtA7942 cAMP:AMP) which is likely to exceed the physiological range of intracellular concentrations by an order of magnitude. The lowest tested cAMP to ADP or AMP ratio of 0.2 (95 µM cAMP: 1.905 mM ADP or AMP) is within physiologically relevant range and cAMP did not affect the association of either SbtA-SbtB pair. Therefore, changes in cAMP concentrations expected to occur *in vivo* are likely inconsequential compared to the variations in intracellular ATP levels and the AEC.

**Fig. 3.**
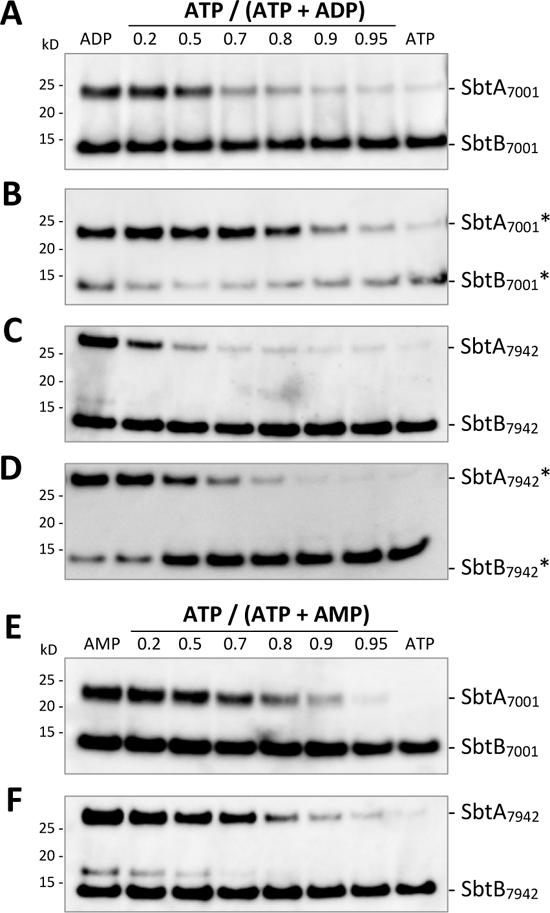
Effects of ATP:ADP and ATP:AMP ratios on the association of SbtA and SbtB proteins. The SbtA_7001_-SbtB_7001_ (**A, C, E**) and SbtA_7942_-SbtB_7942_ (**B, D, F**) pairs were expressed in *E. coli* (**A, C, E, F**) and in *Synechococcus* ΔCS (denoted by *; **B, D**). Variation in energy charge was simulated during *in vitro* IMAC binding assays by different molar ratios of ATP:ADP (**A-D**) and ATP:AMP (**E, F**), with a total adenylnucleotide concentration of 2 mM. Representative immunoblot images of HAHis-tagged SbtB and SbtA proteins retained in SbtA:SbtB complexes of n=3-4 per adenylnucleotide treatment. The 17 kD-protein above SbtB_7942_ is a fragment occasionally detected with the anti-SbtA antibody (**C, F**).

**Fig. 4.**
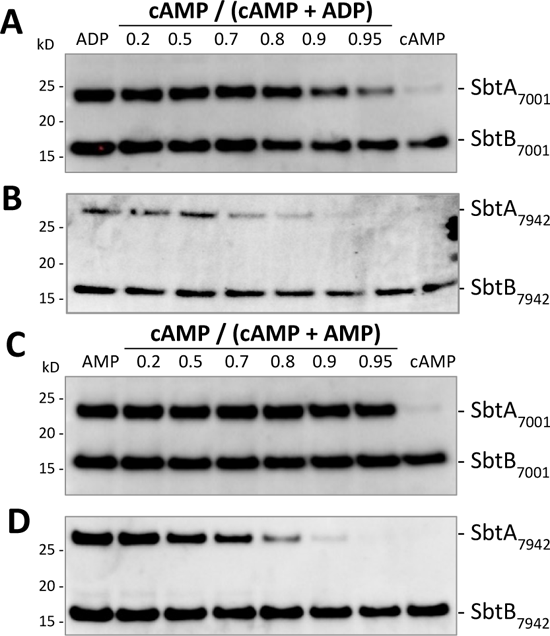
Effects of the cAMP:ADP and cAMP:AMP ratios on the association of SbtA and SbtB proteins. *In vitro* IMAC binding assays were performed across a range of cAMP:ADP (**A, B**) and cAMP:AMP (**C, D**) ratios, with a total adenylnucleotide concentration of 2 mM. The SbtA_7001_-SbtB_7001_ (**A, C**) and SbtA_7942_-SbtB_7942_ (**B, D**) pairs were expressed in *E. coli*. The immunoblot images show HAHis-tagged SbtB and SbtA proteins retained in SbtA:SbtB complexes and are representative of three independent experiments per adenylnucleotide treatment.

**Fig. 5.**
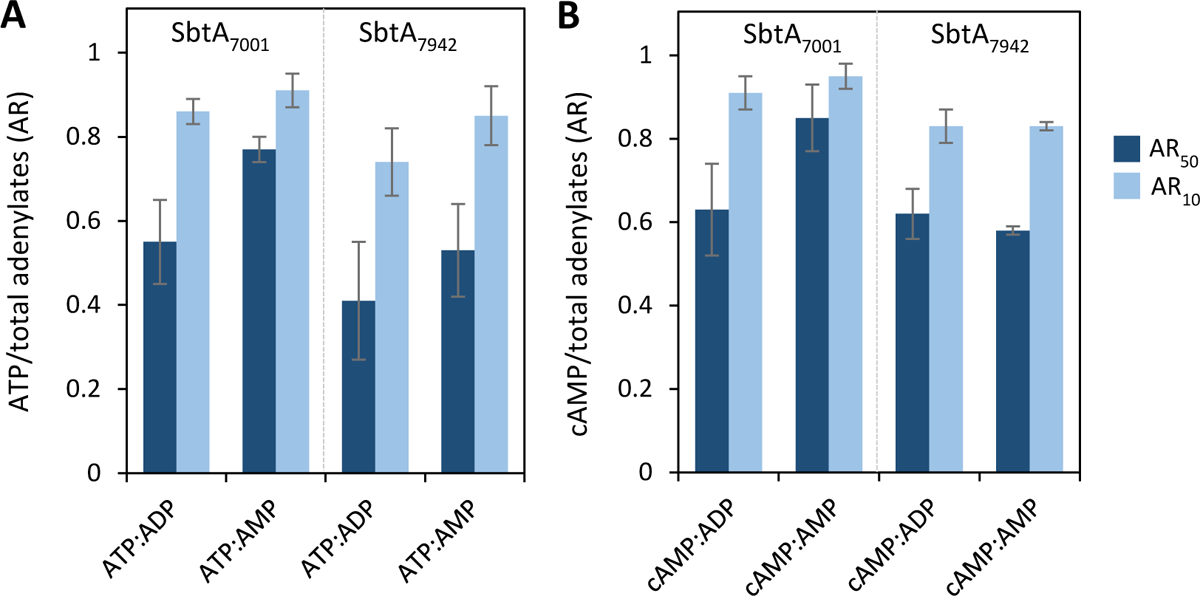
Quantitative estimates for the association of SbtA and SbtB proteins in response to different adenylnucleotide ratios (AR). The ratios of ATP:ADP or AMP (**A**) and cAMP:ADP or AMP (**B**) that supported 50% (AR_50_) and 10% (AR_10_) of SbtA bound to SbtB were estimated from sigmoidal logistic curve fits (Fig. S4). AR values are means (± SE) of 3-4 independent experiments. AR values were not significantly different across each AR category at a *P*=0.05 threshold (AR_50_ *P*=0.232, AR_10_ *P*=0.275).

To compare the association of SbtA with SbtB between adenylnucleotide treatments quantitatively, we estimated the relative changes in SbtA abundance from densitometric analyses of Western blot images (pixel volumes) against ATP:ADP or AMP and cAMP:ADP or AMP ratios (**Fig. S4**). The association of SbtA with SbtB was non-linearly correlated with the adenylnucleotide ratio and fitted best by a sigmoidal logistic model, typical of cooperative ligand binding (mean adjusted R^2^ values of 0.93 ± 0.02; **Fig. S4**). Comparing the adenylnucleotide ratios supporting 50% (AR_50_) or 10% (AR_10_) of the maximum levels of SbtA bound to SbtB, derived from curve fits in **Fig. S4**, differences between the two SbtA-SbtB pairs and in response to the four different adenylnucleotide combinations tested were statistically not significant (**Fig. 5**). However, even though this implied very similar binding affinities of SbtB_7001_ and SbtB_7942_ for those adenylnucleotides as well as comparable ligand-induced binding to the cognitive SbtA proteins, SbtA_7942_ seemed to be dissociated at slightly lower ATP:ADP, cAMP:ADP and cAMP: AMP ratios than SbtA_7001_.

### SbtA activity *in vivo* is modulated by SbtB and the light-stimulated adenylate energy charge

Inside photosynthetic cyanobacterial cells, SbtA and SbtB proteins are both present when Ci supply is low (**Fig. S1A**). In nonphotosynthetic *E. coli*, SbtA appeared to be constitutively inactivated in the presence of SbtB (Du et al., 2014), which implied AEC levels were low under those experimental conditions. In cyanobacteria, however, SbtA is active irrespective of SbtB when photosynthesis is fully light-activated and cells operate at high AEC levels (**Figs. 6, S6, S7**). Therefore, it was of interest to investigate SbtA function in low Ci combined with low AEC states which are prominent during dark-to-low light transitions. Photosynthetic performance and uptake of ^14^C-labelled HCO_3_^-^ in *Synechococcus* ΔCS expressing SbtA_7942_ as the only HCO_3_^-^ uptake system was compared either with or without SbtB_7942_ (SbtAB and SbtA, respectively). Note that concurrent CO_2_ uptake by Ndh-1_3/4_ complexes was eliminated by addition of the carbonic anhydrase inhibitor ethoxyzolamide (EZ), which does not affect SbtA function (**Fig. S5**), and measurements were performed at pH 8.0 where HCO_3_^-^ represents 98% of the Ci pool. Under these conditions, the cells fully depended on SbtA activity for inorganic carbon supply. At light intensities above the light compensation point, O_2_ evolution rates directly reflected the activity of SbtA_7942_. In Ci-starved cell cultures steady-state photosynthetic rates in response to incremental additions of bicarbonate were essentially the same for SbtA and SbtAB with substrate affinities (*K*_0.5Ci_; 9 ± 4 and 30 ± 5 µM Ci, respectively) and maximum rates of photosynthetic O_2_ evolution (*V*_maxO2_; 164 ± 19 and 167 ± 12 µmol mg Chl*a* ^-1^ h ^-1^, respectively) (**Fig. S6**). Similarly, alternating light intensities between 1400 and 200 µmol photons m^-2^ s^-1^ in the presence of saturating Ci levels (250-500 µM) did not lead to discernible differences between SbtA and SbtAB in steady state HCO_3_^-^ uptake or photosynthetic O_2_ evolution rates (**Fig. S7**). Together, this suggested SbtB is not required for and does not regulate SbtA_7942_ activity once photosynthesis has been fully activated in response to light-intensity driven changes in carboxylation rates and Ci demand.

**Fig. 6.**
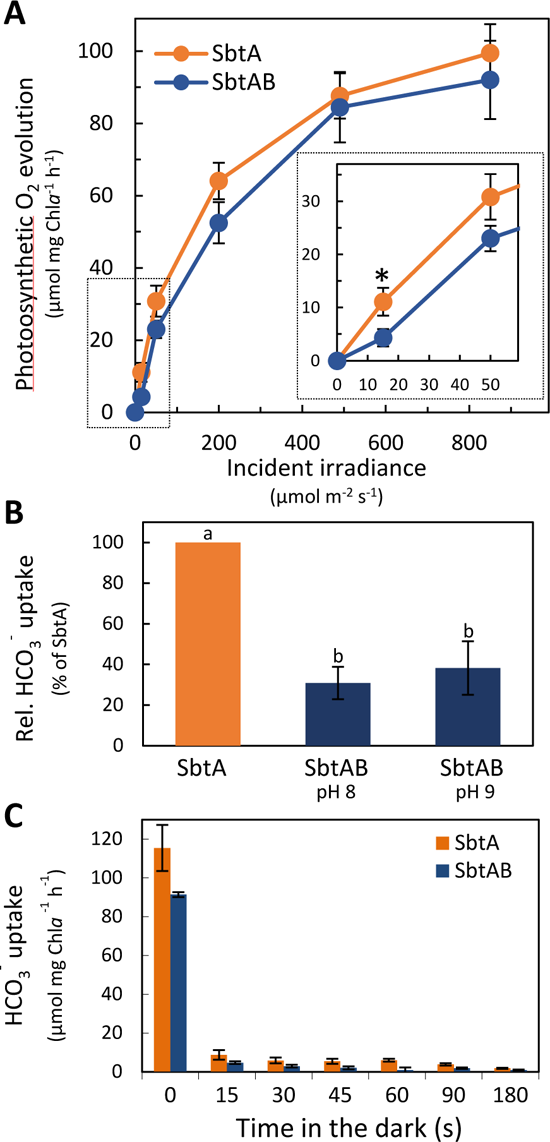
Light-dependent bicarbonate uptake by SbtA is modulated by SbtB in *Synechococcus* ΔCS. The SbtA_7942_ protein was expressed alone (SbtA) or with SbtB_7942_ (SbtAB) in *Synechococcus* ΔCS (ΔCS) at low Ci in the light. CO_2_ uptake was inhibited by EZ to restrict Ci acquisition to bicarbonate uptake by SbtA. (**A**) HCO_3-_ uptake-dependent gross photosynthetic O_2_ evolution rates in response to stepwise increased light intensity after 1 h dark-adaptation. Respiratory O_2_ consumption was the same for SbtA and SbtAB (27 ± 4 and 27 ± 3, respectively). Values are means ± SE; n=4-5. (**B**) Induction of HCO_3-_ transport activity of SbtA in dark-adapted cells immediately after dark-to-low light (30 μmol m^-2^ s^-1^) transitions at pH 8 and pH 9 during 30-60 s pulses of [_14_C]HCO_3-_ uptake, using silicon oil centrifugation-filtration. HCO_3-_ uptake rates are shown relative to cells expressing SbtA alone; n=4-5. Relative HCO_3-_ uptake rates were calculated by subtraction of the unspecific incorporation of _14_C-label, mainly due to extracellular [_14_C] HCO_3-_ in the water shell surrounding each cell and a very small amount of _14_CO_2_ diffusion into the cells. This was derived from _14_C-label incorporation into Ci uptake inhibited ΔCS (38% ± 6 (pH 8) and 42% ± 8 (pH 9). (**C**) HCO_3-_ uptake rates normalized to Chl*a* content at various time points after a light (400 μmol m^-2^ s^-1^) to dark shift. Values are means ± SE; n=3-5. Statistically significant differences (*P* < 0.05) are denoted by an asterisk (**A**) or different letters (**B**).

**Fig 7.**
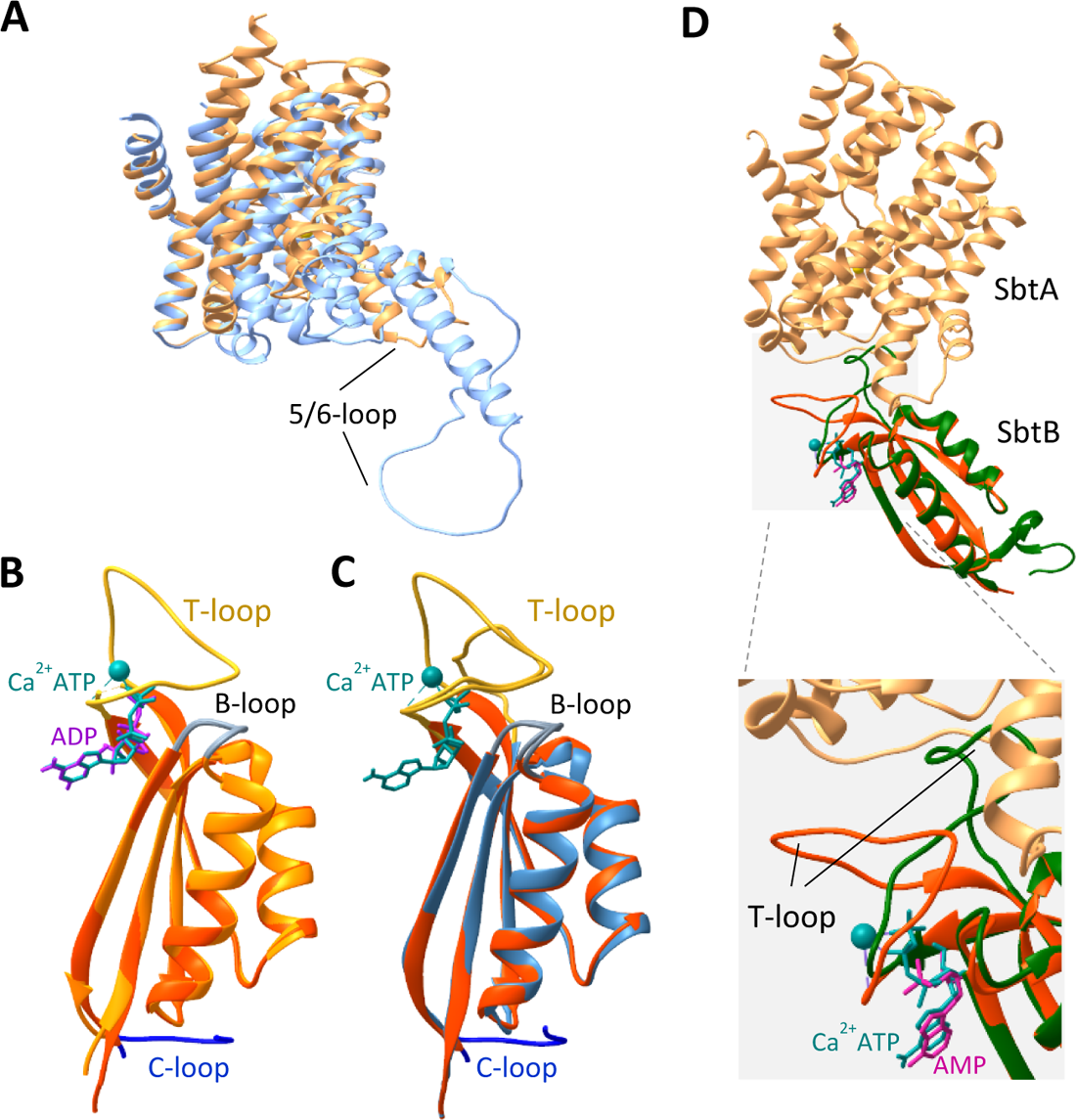
Structural comparison of the overall fold of SbtA and SbtB protein models. (**A**) Superposition of homology models of SbtA_7001_ (light orange) and SbtA_7942_ (light blue). The core domains and arrangement of the transmembrane helices are well matched but the 5/6 loop connecting transmembrane helix 5 and 6 is divergent, protruding further into the cytoplasm on SbtA_7942_. (**B**) Superposition of the three-dimensional models based on crystal structures of SbtB_7001_ with Ca^2+^ and ATP ligands (red-orange, SbtB_7001_-CaATP; PDB: 6MM2) and SbtB_7001_ with ADP bound (orange, SbtB_7001_-ADP; PDB: 6MMC), demonstrating that ligand-induced structural differences affect primarily the T-loop. The T-loop assumes a closed conformation in SbtB_7001_-CaATP and an open, disordered state in SbtB_7001_-ADP which was not resolved in crystals as indicated by the dashed line. (**C**) Superposition the of the homology model of the apo-SbtB_7942_ monomer (blue) on SbtB_7001_-CaATP; both SbtB proteins are predicted to be nearly identical in three-dimensional structures. (**D**) Superposition of the SbtA_7001_ and SbtB_7001_-CaATP onto a SbtA_6803_ monomer (PDB: 7EGK, chain C) associated with a SbtB680-AMP monomer (chain D) which were extracted from the trimeric SbtAB_6803_ cryo-EM structure (PDB: 7EGK). Since the three-dimensional architecture of SbtA_7001_ and SbtA_6803_ are highly similar (Fig. S12), only SbtA_7001_ is shown for clarity. The core structure of SbtB_7001_-CaATP and SbtB_6803_-AMP is nearly identical, but the T-loop conformations diverge in a ligand-specific manner consistent with the observed SbtA:SbtB interaction in this study (enlarged). Binding of AMP to SbtB_6803_ (green) promotes an open T-loop conformation which extends into SbtA consistent with SbtA:SbtB association, whereas Ca_2+_ATP bound to SbtB_7001_ (red-orange) promotes folding of the T-loop away from SbtA.

In contrast, SbtB clearly modulated SbtA function in the initial phase of dark-to-low light transitions. After an hour of dark-adaptation, cells co-expressing SbtA and SbtB were unable to activate SbtA_7942_-dependent photosynthetic O_2_ evolution to the same extent as cells expressing SbtA alone during the initially low light phase of light response measurements, when light levels were below or near the light compensation point of 10-15 µmol photons m^-2^ s^-1^ (**Fig. 6A**). Low light activation of ^14^C-labelled HCO ^-^ uptake by SbtA_7942_ clearly showed SbtA activation was strongly suppressed in the presence of SbtB_7942_ during the first 30-60 s after long-term dark-adaptation, reducing net SbtA activity to 31% and 38% of the activity observed in the absence of SbtB at pH 8 and pH 9, respectively (**Fig. 6B**). Since many photosynthetic processes are almost instantaneously inactivated in the dark, we tested whether SbtB affected [^14^C]HCO_3_ ^-^ uptake by SbtA during light to dark transition. Interestingly, SbtA_7942_ was fully inactivated within 15 s after cells were shifted from light into the dark without any effect on the inactivation kinetics by the presence of the SbtB_7942_ protein (**Fig. 6C**). Altogether this suggests, SbtA activity is regulated at multiple levels, which may be SbtB-independent, but also involve SbtB and AEC coupled regulation of SbtA activity demonstrated by dark-to-light activation kinetics of SbtA.

In *Synechocystis sp.* PCC6803, deletion of SbtB_6803_ was shown to reduce growth under low Ci condition in low light (Selim et al., 2018), and a broader regulatory function of CCM components by SbtB was proposed. If this was the case in *Synechococcus* ΔCS, one would expect a negative effect on growth in cells expressing SbtA alone compared to those co-expressing SbtB and SbtA. However, photoautotrophic growth of cells expressing either SbtA_7942_ or both SbtA_7942_and SbtB_7942_, was very similar when supplied with air levels of CO_2_ under diurnal light-dark cycles (**Fig. S8**), suggesting deletion of SbtB_7942_, unlike SbtB_6803_, did not impair photoautotrophic growth of *Synechococcus* ΔCS under the conditions that provide a relatively stable light environment and sufficient CO_2_ and nutrient supply.

## Discussion

Allosteric regulation of cyanobacterial bicarbonate transporters has been an unresolved question for quite some time with patchy evidence for diverse mechanisms and co-factors that may be involved. It has already been established that SbtB is capable of exerting negative regulation on SbtA bicarbonate uptake activity in a heterologous expression system (Du et al., 2014). Here, we provide evidence for regulation of two SbtA bicarbonate transporters by protein-protein interaction with their cognate PII-like SbtB proteins in response to the *in vitro* simulation and *in vivo* modulation of the adenylate energy charge. The SbtA-SbtB pairs originated from the transitional α-cyanobacterium *Cyanobium sp.* PCC7001 and the ẞ-cyanobacterium *Synechococcus elongatus* PCC7942, which are phylogenetically relatively distant species (Rae et al., 2011; Sánchez-Baracaldo et al., 2019) but share overall sequence homology at the amino acid level, with 60% identity between the two SbtA (**Fig. S9A, S10**) and 58% identity between the two SbtB (**Fig. S9B, S11**) proteins. Importantly, the modelled structures of SbtA_7001_ compared to SbtA_7942_ and SbtB_7001_ compared to SbtB_7942_ are nearly identical based on three-dimensional models (**Fig. 7**). Together with convincingly similar interaction profiles for both SbtA-SbtB pairs, we concluded they likely share the same overall regulatory mechanisms. It should be noted that about 50% of the cyanobacterial SbtB family, which includes the SbtB proteins studied here, lack a putative redox regulatory domain on the C-terminus, as typically found on SbtB of *Synechocystis sp.* PCC6803 (Selim et al., 2023). This makes the present two SbtAB pairs a simpler system for studying the nature of the primary signal(s) that direct allosteric inactivation/activation of SbtA.

### The AEC rather than cAMP is the probable primary effector determining SbtA:SbtB interactions

Intracellular adenylnucleotide pools invariably contain ATP, ADP, AMP and cAMP, albeit relative concentrations may vary rapidly dependent on metabolic activity in response to environmental and developmental factors (De la Fuente et al., 2014). In cyanobacteria, cAMP is predominantly a secondary messenger participating in different signalling events, whereas the relative abundances of ATP, ADP and AMP define and communicate the cellular energy status. Notwithstanding, both processes could translate Ci availability and Ci demand into SbtB-mediated regulation of SbtA. In addition, relative abundances of cAMP, ATP and AMP in the adenylate pool are interdependently influenced by the activities of adenylate cyclase and phosphodiesterases which catalyze cAMP synthesis from ATP and hydrolytic degradation to AMP, respectively; depending on the species, adenylate cyclase activity is stimulated by N and P deficiency, osmotic stress, light quality and in some cases also bicarbonate or CO_2_ (Cann, 2004; Hammer et al., 2006; Xu and Su, 2009; Agostoni and Montgomery, 2014), potentially linking cAMP and Ci sensing.

Initially, evidence from *Synechocystis sp.* PCC6803 suggested the cAMP:AMP ratios determined formation of SbtA:SbtB complexes and SbtA activity (Selim et al., 2018). SbtB_6803_ was proposed to be a novel cAMP receptor sensing the Ci status and mediating acclimation of the CCM to high Ci levels, based on impaired acclimation in mutant cell lines lacking SbtB. In that study, high Ci acclimated cells showed the expected general downregulation of Ci uptake alongside relatively higher cAMP levels and cAMP:AMP ratios than in low Ci acclimated cells. However, SbtB_6803_ was dissociated from SbtA_6803_ which implied SbtA was inactive in its unbound state. Conversely, bicarbonate uptake was upregulated in low Ci acclimated cells with high AMP levels and low cAMP:AMP ratios, which favoured formation of SbtA:SbtB complexes, and therefore suggested SbtA was active when bound to SbtB. Those findings were difficult to reconcile with the simple interaction model for SbtB-mediated SbtA regulation in *E. coli*, where SbtA alone showed activity and co-expression of SbtA and SbtB inhibited SbtA-dependent bicarbonate uptake, including the SbtA_6803_-SbtB_6803_ pair (Du et al., 2014). Meanwhile, regulation of SbtA_6803_ has been revised based on the finding that SbtB_6803_ displays high affinity binding to SbtA only with AMP as ligand and when the R-loop Cys residues are oxidized, which appeared to be the only condition for T-loop conformation that permits binding to SbtA (Selim et al., 2023).

The binding analyses performed here (**Fig. 3**), and the fact that the R-loop is absent from SbtB_7001_ and SbtB_7942_ (**Fig. S9B**), clearly support the view that low levels of ATP and low ATP:ADP and AMP ratios, representing a low AEC, determine the association of SbtA_7001_-SbtB_7001_ and SbtA_7942_-SbtB_7942_ pairs rather than cAMP levels and redox state of SbtB. Considering, that the cAMP concentrations that effectively prevented the *in vitro* association of SbtA and SbtB were up to 1000-fold higher than physiological levels of CAMP in cyanobacteria (Ohmori and Okamoto, 2004), it seems unlikely the intracellular concentration of cAMP required for effective competition with ATP for binding sites on SbtB would occur *in vivo*. Furthermore, the 6-12 times higher binding affinity of SbtB_7001_ for ATP over cAMP, ADP or AMP (Kaczmarski et al., 2019) is consistent with the proposed AEC-dependent regulation of these SbtA-SbtB pairs. This AEC control of SbtA via binding/unbinding of SbtB might be a feature across many cyanobacteria, but further experimentation is required to confirm this.

### The SbtA and SbtB proteins interact dynamically in response to the adenylate energy charge

ATP, ADP and AMP form the core of the energy system of living cells. The majority of biochemically available energy is linearly correlated with the molar fraction of ATP and ½ molar fraction of ADP of the total adenylate pool, but was experimentally simplified here by testing effects of ATP:ADP and ATP:AMP ratio effects separately (**Fig. 3**). Adenylate pools measured from a diverse range of tissues, organisms and environmental conditions suggest active growth requires an adenylate energy charge between 0.8 and 0.9, while a drop of the AEC below 0.5 is unsustainable (Chapman et al., 1971). Under favourable environmental conditions, the AEC is kept remarkably steady near 0.8 in cyanobacteria the light (Rust et al., 2011). Regulatory proteins often sense the AEC through competitive binding of ATP or ADP (Atkinson and Walton, 1967), which includes PII proteins that regulate target enzyme activity by undergoing specific conformational changes in response to binding ATP or ADP, which alters the interaction between the PII protein and its target and modulates the activity of the target protein (Radchenko et al., 2013; Zeth et al., 2014; Selim et al., 2019). This principle also appears to apply to the interaction of the PII-like SbtB proteins with respective SbtA transporters from both cyanobacterial species investigated. More than 90% of SbtA was unbound (proposed active state) at a simulated AEC of 0.74 to 0.91 (**Figs. 3, 5**) approximating measured ATP/(ATP+ADP) ratios of 0.8 to 0.9 reported for *Synechococcus elongatus* (formerly *Anacystis nidulans*) ((Rust et al., 2011), recalculated from (Bornefeld and Simonis, 1974)) and the thermophile *Synechococcus* strain Y-7c-s in medium light (250-270 µmol m^-2^ s^-1^) (Kallas and Castenholz, 1982). This suggests, in the light, photosynthetically active cyanobacterial cells would maintain an almost fully active pool of SbtA, if the AEC was the defining parameter. The fact that SbtA_7942_ activity was the same for photosynthetically fully activated *Synechococcus* ΔCS strains expressing either SbtA or SbtA and SbtB (**Fig. S6**, **S7**) supports the notion that SbtB had in fact fully dissociated from SbtA and did not limit SbtA activity at high AEC. Without photosynthetic ATP production in the dark, or during excessive cellular energy demand such as sudden decreases in light intensity and nutrient, pH or temperature stress, the AEC is likely to decrease substantially in cyanobacterial cells. In *S. elongatus*, the ATP:ADP ratio, was reduced to 0.55-0.6 about 10 min after light to dark transfer (Kallas and Castenholz, 1982; Rust et al., 2011). Based on our *in vitro* binding analysis, this suggests, on average more than 50% of both SbtA7001 and SbtA7942 should be bound to their respective SbtB proteins and form a stably-inactivated SbtA pool in prolonged darkness. During light exposure, after long dark periods, it can take from seconds to minutes for the cellular AEC to reach 0.8-0.9, dependent on light intensity and activation of CO_2_ fixation by Rubisco and the CBB cycle enzymes. Indeed, low light (70-100 µmol m^-2^ s^-1^) did not increase the AEC beyond 0.6 in thermophilic *Synechococcus* (Kallas and Castenholz, 1982).

Importantly, when *Synechococcus* ΔCS strains expressing either SbtA or SbtA and SbtB were analysed in dark-to-low light transitions, equivalent to raising AEC ratios from about 0.5 to 0.8-0.9, we detected a transient and marked retardation in SbtA-dependent bicarbonate uptake, decreasing SbtA activity by 60-70% when SbtB was present compared to SbtA alone in the initial low light (30-50 µmol m^-2^ s^-1^) phase (**Fig. 6A, B**). Assuming an AEC of 0.6 in low light, similar to Kallas *et al*., around 50% of SbtA should still have been associated with SbtB (**Figs. 3, 5**). In our view, this is a convincing indication that a rise in AEC is needed to displace the inhibitory SbtB curfew protein for SbtA activation during this transition and that this takes a few seconds to achieve. Despite the technical difficulty, future work looking at correlations between time-resolved AEC changes and bicarbonate uptake in *Synechococcus* ΔCS strains should be able to test this further.

### Adenylnucleotide-dependent conformation changes of SbtB aligns with regulatory binding to SbtA

Structural information of SbtA:SbtB complexes and binding of adenylate ligands to SbtB strongly support the aforementioned AEC-dependent regulation of the SbtA:SbtB interaction *in vitro*. Inevitably, uncertainty exists in extrapolating from *in vitro* findings to implications for *in vivo* regulation. However, the case of regulation of SbtA by adenylate charge is at least feasible, especially for the significant class of SbtA-SbtB-containing species that lack the putative C-terminal redox regulatory domain on SbtB. The core structures and key amino acids involved in ligand binding that are thought to facilitate SbtA:SbtB interactions are highly conserved between crystal structures of SbtB_6803_ and SbtB_7001_ and homology models of SbtB_7942_ (**Figs. 7B,C, S10**) as well as crystallized SbtA_6803_ and homology models of SbtA_7001_ and SbtA_7942_ (**Figs. 7A,C, S10A, S12**), which lends some validity to propose a generalized model for the regulatory interaction of subsets of those SbtB proteins missing the R-loop and their cognate SbtA proteins. In PII proteins, the surface-exposed, flexible T-loop has been shown to mediate the regulatory interaction with target protein(s). Upon binding small effector molecules, ligand-induced conformation changes determine whether PII binds to its target. For example, binding of ATP to the PII protein from *Synechococcus elongatus* PCC7942 induced T-loop conformation changes that facilitated association with N-acetyl-glutamate kinase (NAGK) and arginine synthesis (Zeth et al., 2014; Selim et al., 2019), and the ADP-stabilized T-loop of the PII protein GlnK from *E. coli* enable formation of an inactivating complex with the ammonium transporter AMTB (Radchenko et al., 2010). Crystallographic data support a direct correlation, analogous to PII proteins, between adenylate-ligand induced changes in T-loop conformation of SbtB proteins (Selim et al., 2018; Kaczmarski et al., 2019; Fang et al., 2021; Selim et al., 2023) and can explain the opposite effects of ATP and ADP or AMP on SbtA:SbtB binding described here. The probability for each adenylate ligand to occupy a nucleotide binding site would depend on the binding affinity and the local concentration of the ligand. At high ATP:ADP or AMP ratios, when predominantly Ca^2+^ATP is likely to occupy the nucleotide binding sites of the SbtB_7001_ trimers, the T-loops are probably stabilized (**Fig. 7B,C**) through interactions of specific residues with the γ-phosphate of ATP, in a position close to the core overlapping the binding site (Kaczmarski et al., 2019). When superimposed onto SbtB_6803_ in the trimeric SbtA-SbtB cryo-EM structure, it becomes apparent that the Ca^2+^ATP-induced stable T-loop conformation of SbtB_7001_ interferes with binding to SbtA (**Fig. 7D**), which would allow SbtA to be fully active. In contrast, at low ATP:ADP or AMP ratios, occupation of the binding site mostly by ADP or AMP (**Fig. 7B,C**) would leave the T-loops in a disordered or extended conformation (Kaczmarski et al., 2019; Fang et al., 2021), which would enable formation of inactivated SbtA:SbtB complexes.

### A proposed model implicating SbtB in the regulation of SbtA activity in cyanobacteria

For cyanobacteria, it is a selective advantage to maximize photosynthetic CO_2_ fixation and carbon gain for growth in the light. In the dark, photosynthetic CO_2_ fixation ceases making Ci acquisition redundant and energetically futile. Additionally, continued accumulation of large amounts of bicarbonate may disrupt intracellular pH and ion homeostasis, particularly if [Na^+^] were to build up by HCO_3_^-^/Na^+^ symport due to SbtA activity. Therefore, the co-ordinated regulation of Ci acquisition and photosynthetic activity may be an imperative for optimal survival and success of cyanobacteria. A hypothetical scenario for the regulation of bicarbonate uptake by SbtA in cyanobacteria is depicted in **Fig. 8**, assuming functional units of both SbtA and SbtB proteins are trimers as indicated by cryo-EM and crystal structures (Kaczmarski et al., 2019; Fang et al., 2021). In *Synechococcus*, SbtA function is subject to several regulatory mechanisms operating at different time scales. In constant light and stress-free environments, bicarbonate uptake rates of SbtA are clearly a product of light intensity and Ci availability, independent of the SbtB protein (**Figs. 6**, **S7**). In light/dark cycles, SbtA is turned on and off within a few seconds (**Fig. 6C**) in sync with the light-induced changes in Ci demand for photosynthetic carboxylation, likely preventing the aforementioned complications associated with bicarbonate and Na^+^ over-accumulation. The nature of the signals and signal transduction pathways triggering the fast activation-deactivation kinetics of SbtA may be complex and cannot be precisely known at this stage, but some details can be envisaged. SbtA-mediated bicarbonate uptake is immediately inactivated after a light-dark transition supposedly associated with a drop in the AEC, assuming a similar decrease of the ATP/(ATP+ADP) ratio from 0.85 to about 0.4 observed in *S. elongatus* previously (Bornefeld and Simonis, 1974; Takano et al., 2015). If AEC-induced binding of SbtB was the primary inactivation mechanism during the light-to-dark transition, one would expect de-regulation of SbtA activity in the absence of SbtB. Yet, SbtA-mediated bicarbonate uptake ceased within 15 s in the dark irrespective of the presence of SbtB (**Fig. 6C**), which suggested the initial fast inactivation of SbtA may be mediated by any one or a combination of other signals derived from the many dark-induced changes in the cell which may involve the more oxidized redox state (Tamoi et al., 2005), changes in net availability of ATP, in plasma membrane energization potentially involving Ca^2+^ signalling (Torrecilla et al., 2004), in the transmembrane Na^+^ gradient, and in protein post-translational modifications such as phosphorylation (Spät et al., 2015; Angeleri et al., 2016). Many of these changes are likely faster than a possible low-AEC-induced SbtA:SbtB association. On the other hand, we did detect significant differences in the activation of bicarbonate uptake upon transition from dark to light (Fig. 6), indicating low AEC-induced formation of inactive SbtA:SbtB complexes in the dark delayed activation of SbtA until light-dependent increase in AEC was sufficient to facilitate the dissociation of SbtB and SbtA.

**Fig. 8.**
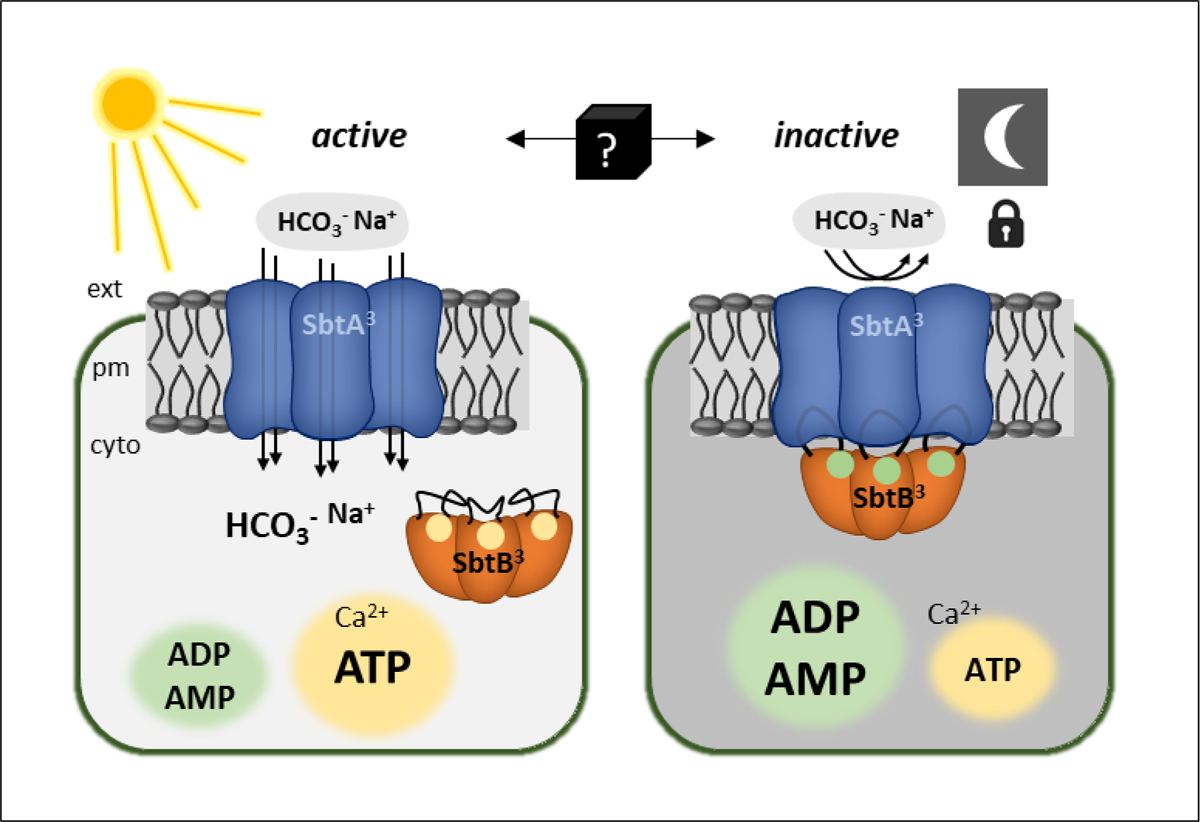
Proposed model for regulation of SbtA bicarbonate transport activity involving SbtB by environmental stimuli and the adenylate energy charge (AEC). In the light, expression of SbtA and SbtB in *Synechococcus elongatus* is induced by low Ci availability. The plasma membrane (pm)-integral SbtA trimer(SbtA_3_) actively transports bicarbonate from the external environment (ext) into the cytoplasm (cyto) presumably dissociated from SbtB. SbtA activity depends on Na^+^-symport and requires a cell-inward-directed Na^+^ gradient with a probable HCO_3-_:Na^+^ symport stoichiometry of 1:1 through each monomer. Light-stimulated photosynthetic electron transport sustains high levels of intracellular ATP over ADP and AMP, i.e. high AEC, which facilitates binding of Ca^2+^ATP to SbtB trimers (SbtB_3_), stabilizing the SbtB T-loops, and SbtB remains dissociated from SbtA. In prolonged darkness, cells operate at a lower AEC. Relatively higher amounts of ADP and AMP replace ATP as ligands in SbtB trimers, which favours open T-loops, and inactive SbtA:SbtB complexes are formed. In such scenario, SbtA reactivation after a transition from dark to light would depend on the rate of increase in AEC and concomitant AEC-dependent dissociation of SbtB.

## Conclusion

The adenylate-specific binding of SbtB to SbtA suggests that *in vivo* SbtA trimers become locked into an inactive state when the cellular adenylate energy charge is low, as is the case in low light or darkness. This becomes evident from delay of SbtB-dependent re-activation of SbtA. According to our hypothesis, low AEC-induced binding of SbtB to SbtA provides a curfew mechanism that ensures SbtA activity is tuned to the capacity for photosynthetic C-fixation and tied to the energy resources available. This mechanism may have evolved to override environmental and intrinsic cues that may otherwise induce futile activation/inactivation cycles or activation of SbtA under conditions that demand strategic investment of energy in cell survival rather than maximum growth. Limited light, prolonged nutrient stress or environmental stress may require channelling of resources into maintenance and protective processes instead of maximum photosynthesis and Ci accumulation. It will be of interest to investigate to which extent AEC/SbtB-mediated regulation SbtA has been adopted across cyanobacterial species. The general mechanism of energy charge-sensing via SbtB and its interaction with SbtA is increasingly supported by experimental evidence, however, the precise details are far from being fully understood. Further elucidation of molecular and biochemical mechanisms and further substantiation of the role of the AEC in SbtA:SbtB complex formation *in vivo* under different environmental conditions are challenges for future work. Nonetheless, the link between SbtB as a sensor of the cellular energy status and response regulator of SbtA activity is an important novel insight which may aid development of informed strategies for expression of SbtA-SbtB systems in higher plant chloroplasts.

## Materials and Methods

### Bacterial and cyanobacterial cell lines

Nucleotide and amino acid sequences were retrieved from the Integrated Microbial Genome database (IMG; https://img.jgi.doe.gov) and are referred to by gene ID and Locus_tags. The commercially available *Escherichia coli* strain DH5⍺ (Invitrogen) was used for cloning and heterologous expression of SbtA and SbtB proteins for binding studies. The HCO_3_^-^ uptake deficient cell line *Synechococcus* ΔCS was derived from wild-type *Synechococcus elongatus* PCC7942 by sequential insertional inactivation of the only two endogenous bicarbonate transporters, BCT1 (*cmpABCD* operon) and SbtA (*sbtAB* operon). BCT1 expression was eliminated by replacement of the *cmpA* (IMG ID: 637799921, Synpcc7942_1488) and part of the *cmpB* (IMG ID: 637799922, Synpcc7942_1489) genes with a selective kanamycin resistance (*kan^R^*) cassette (Omata et al., 1990). To this end, a 2.5 kb genomic DNA fragment spanning the *cmpA* and *cmpB* genes was amplified by PCR. The *kan*^R^ marker was inserted into the central *Bgl*II site of the *cmpAB* fragment, generating gene-specific 3’ and 5’ flanking sequences for homologous recombination (**Table 1**) and ligated into pUC18 (pUC18::Δ*cmpAB*) which is a non-replicating vector in cyanobacteria. The pUC18::Δ*cmpAB* plasmid was transformed into wild-type and generated the single deletion mutant (ΔC). For the second deletion (ΔS), the full length *sbtA* (IMG ID: 637799907, Synpcc7942_1475) and *sbtB* (IMG ID: 637799908, Synpcc7942_1476) operon was replaced with a chloramphenicol resistance (*cm^R^*) marker gene (Dzelzkalns et al., 1984) in *Synechococcus* ΔC. Two 1100 and 1400 bp genomic DNA fragments up- and downstream of the *sbtAB* operon, respectively, were amplified by PCR (**Table 1**), attached as 3’ and 5’ genomic flanking sequence to the *cm^R^*marker gene and inserted into plasmid pUC18 (pUC18::Δ*sbtAB*), which was then transformed into *Synechococcus* ΔC to generate the double deletion strain (ΔCS). Cyanobacterial transformation was carried out by natural DNA uptake as described previously (Golden and Sherman, 1984). Single cell-derived *kan^R^* and/or *cm^R^* colonies were selected on the appropriate antibiotic-containing media (kanamycin 100 µg ml^-1^, chloramphenicol 8 µg ml^-1^) until ΔC and ΔS deletions had fully segregated, which was verified by PCR using primer pairs Cmp-F/Cmp-R and SbtA-F/SbtB-R, respectively (**Table 1**).

**Table 1.**
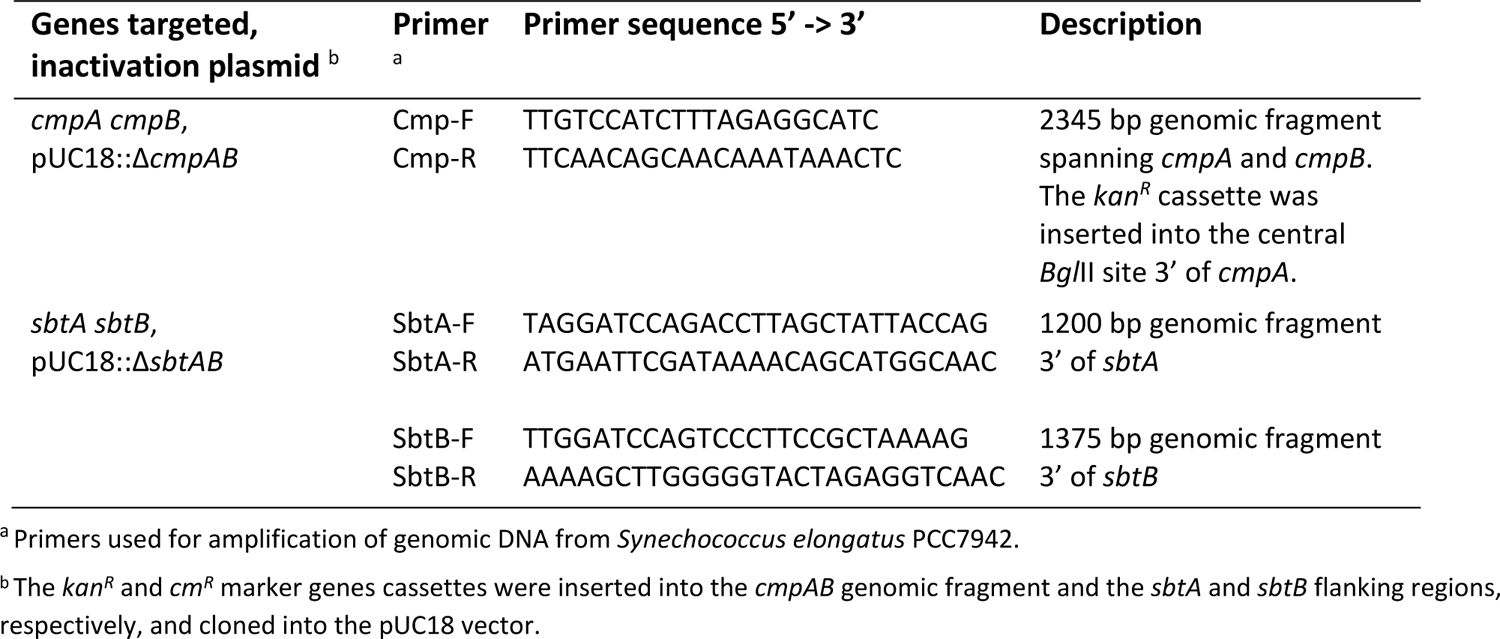
Primers for generation of the bicarbonate transporter-deficient *Synechococcus* ΔCS mutant.

For protein expression, *sbtA* and *sbtB* genes from *Synechococcus elongatus* PCC7942 and *Cyanobium sp.* PCC7001 (IMG ID: 647590134, CPCC7001_1784 and IMG ID: 647590133, CPCC7001_1671, respectively) were synthesized (GenScript, New Jersey, USA) and inserted into the *Sac*I/*Xba*I restriction sites of plasmids pSE2-1 and pSE4-1 which both carry a spectinomycin resistance (*sp^R^*) marker gene (Price et al., 2004; Du et al., 2014) and contain origins of replications for maintenance in *E. coli* and in *Synechococcus elongatus*. Genes of interest were expressed from pSE2-1 under control of the isopropyl β-D-1-thiogalactopyranoside (IPTG)-inducible *lacZ* promoter and *lacIQ* repressor and from pSE4-1 using their native *sbtA* and *sbtB* promoters. The SbtB proteins were C-terminally tagged with a HA epitope (YPYDVPDYA) for immunodetection and 5 or 6 His residues for affinity purification in IMAC assays. For expression of SbtA alone, the appropriate *sbtB* genes were removed by restriction digestion and re-ligation from the *sbtAB* gene inserts in pSE2-1 and pSE4-1. Transformed *E. coli* and cyanobacterial cells were selected on solid media containing 100 and 10 µg ml^-1^ spectinomycin, respectively. Cell lines and expression vectors used in this study are listed in **Table 2**.

**Table 2.** *E. coli* and cyanobacterial cell lines and plasmids used for protein expression and physiological analyses of the SbtA and SbtB proteins from *Cyanobium sp.* PCC7001 and *Synechococcus elongatus* PCC7942.

| Organism | Plasmid::Gene(s) of interest<sup>a</sup> | Promoter(s) | Proteins expressed |
| --- | --- | --- | --- |
| *E. coli* DH5α | pSE2-1::sbtAsbtBH6-7942 | *lacZ* | SbtA<sub>7942</sub><br>SbtB<sub>7942</sub>-HAHis |
| *E. coli* DH5α | pSE2-1::sbtAsbtBH5-7001 | *lacZ* | SbtA<sub>7001</sub><br>SbtB<sub>7001</sub>-HAHis |
| *Synechococcus* ΔCS | pSE4-1::sbtA-7942 | *native sbtAB-7942*<sup>b</sup> | SbtA<sub>7942</sub> |
| *Synechococcus* ΔCS | pSE4-1::sbtAsbtB-7942 | *native sbtAB-7942*<sup>b</sup> | SbtA<sub>7942</sub><br>SbtB<sub>7942</sub> |
| *Synechococcus* ΔCS | pSE4-1::sbtAsbtBH6-7942 | *native sbtAB-7942*<sup>b</sup> | SbtA<sub>7942</sub><br>SbtB<sub>7942</sub>-HAHis |
| *Synechococcus* ΔCS | pSE4-1::sbtAsbtBH6-7001 | *native sbtAB-7001*<sup>c</sup> | SbtA<sub>7001</sub><br>SbtB<sub>7001</sub>-HAHis |

<sup>a</sup> pSE2-1 and pSE4-1 contain a spectinomycin resistance cassette (*sp<sup>R</sup>*) for selection.
<sup>b</sup> *sbtA-7942* and *sbtB-7942* are transcribed in opposite directions away from each other. The intergenic region between the ATG-translation start codons contains used for expression contains the promoters both genes.
<sup>c</sup> *sbtA-7001* and *sbtB-7001* are transcribed in the same direction in tandem. The promoter is contained within the 500 bp upstream of the GTG-translation start for *sbtA*.

### Growth of *E. coli* and cyanobacteria

For maintenance, mutant selection and protein expression, *E. coli* DH5α derived cell lines were grown at 37 °C in Lysogeny broth (LB) supplemented with 100 µg ml^-1^ spectinomycin as described previously (Du et al., 2014), and expression of SbtA and SbtB proteins was induced for 3 h with 1 mM IPTG. *Synechococcus* ΔCS cell lines were grown at 30 °C in modified BG-11 medium (buffered with 20 mM HEPES-KOH pH 8.0 (Maeda et al., 2002; Woodger et al., 2003) in high Ci (2-4% (v/v) CO_2_ enriched air) or low Ci (ambient air, 400 ppm CO_2_) under constant light (70 µmol m^-2^ s^-1^). Antibiotics were supplied as appropriate (100 µg ml^-1^ kanamycin, 8 µg ml^-1^ chloramphenicol, 100 µg ml^-1^ spectinomycin). Protein expression of SbtA and SbtB was induced by exposing high Ci-grown cyanobacterial cultures for 4 h to low Ci. For growth assays, dilution series of low Ci-induced cultures grown to mid-log phase were adjusted to OD_730_ of 0.2 and a dilution series spotted onto agar plates containing growth medium with antibiotic supplements. Plates were then incubated in air under 50 µmol m^-2^ s^-1^ white light (photoperiod, 12 h light: 12 h dark) for 10 days. Final biomass accumulation was recorded as scanned images.

### Extraction of native proteins

The method for the isolation of membrane-enriched fractions of native proteins from *E. coli* and *Synechococcus* ΔCS was adapted from previous work (Du et al., 2014). After induction of protein expression, cultures of *E. coli* or S *Synechococcus* ΔCS were harvested in mid-log phase by centrifugation (5 min, 6000 *g*, 23 °C). Cell pellets were washed once with HEPES lysis buffer (50 mM HEPES, 100 mM NaCl, 5% (v/v) glycerol) and treated with 300 U/µl r-Lysozyme (Sigma, USA) for 15 min at 23 °C. Cells were broken with 0.1 mm Zirconia-silica beads (Biospec Products) in a mini-beadbeater 16 (Biospec Products) at 3450 oscillations min^-1^ for 6 min in HEPES IMAC binding buffer (50 mM HEPES, 100 mM NaCl, 25 mM imidazole, pH 8.0) and 0.1% (v/v) protease inhibitor cocktail (Sigma, USA) and centrifuged for 30 s at 16000 *g* to precipitate cell debris and beads at room temperature. The supernatant was then centrifuged (15 min, 16000 *g*, 4 °C) to pellet the membrane-enriched fraction. Membrane proteins were solubilized in IMAC HEPES binding buffer (50 mM HEPES, 100 mM NaCl, 25 M imidazole, pH 8.0) supplemented with 1% (w/v) *n*-dodecyl β-D-maltoside (DDM) by gentle agitation on a rotator for 30 min at 23 °C, and insoluble matter precipitated by centrifugation (5 min, 16000 *g*, 4°C) prior to further analysis.

### Immobilized metal affinity chromatography (IMAC) protein binding assays

The association of SbtA and HAHis-tagged SbtB proteins was analysed in IMAC binding assays adapted from previous work (Du et al., 2014). Native protein extracts from *E.coli* or *Synechococcus* ΔCS were incubated in IMAC HEPES binding buffer supplemented with potential effector molecules and Ni-charged iminodiacetic acid Profinity™ IMAC Resin (Bio-Rad, USA) on a low speed rotator for 1 h at 23°C. This allowed the His-tag on SbtB to bind to the resin and SbtA to associate with the immobilized SbtB protein. The resin had been prewashed three times with IMAC HEPES binding buffer, and one volume of resin was added to four volumes of the reaction mixture. Depending on the experiment, varying concentrations of MgCl_2_, CaCl_2_, NaHCO_3_, Na_2_HPO_4_, RUBP, 3PGA, 2OG, ATP, ADP, AMP and cAMP were added to the protein extract-resin mixture as potential effector molecules. Standard binding assays were performed with 200 nM CaCl_2_ unless stated otherwise. Further details on effector molecules and adenylnucleotide ratios are provided in the figures and captions. The reaction mixture was transferred onto columns, washed with five volumes IMAC HEPES binding buffer by gravity flow through, which removed unbound protein and effector molecules. SbtB was then eluted from the resin with IMAC HEPES elution buffer (50 mM HEPES, 300 mM NaCl, 250 mM imidazole, pH 8.0). The eluted fraction contained the proportion of SbtB protein immobilized on the beads and the amount of SbtA protein retained due to binding to SbtB. IMAC purified SbtA:SbtB complexes were subjected to immunoblot analysis.

### Immunodetection of SbtA and SbtB proteins and quantification of SbtA:SbtB binding

Aliquots of membrane-enriched and eluted IMAC fractions were supplemented with 2.5% (v/v) β-mercaptoethanol and sample loading buffer (62.5 mM Tris pH 8.0, 2% SDS, 10% glycerol, 0.02% (w/v) Coomassie G250, 0.0025% (w/v) Phenol red), heated for 3 min at 80 °C, centrifuged (30 s, 12000 *g*, 23°C) and separated on linear gradient 4-12% Bis-Tris polyacrylamide gels (NuPAGE® Novex®, Invitrogen, USA) in MES running buffer (50 mM MES, 50 mM Tris base, 0.2% SDS, 1 mM EDTA free acid). Proteins were transferred onto 0.45 μm PVDF (polyvinylidene fluoride) membrane using a semi-dry blotter (Bio-Rad, USA) and Tris-Glycine transfer buffer (48 mM Tris base, 38 mM glycine, 0.038% SDS, 20% methanol). Immunological detection was performed in Tris-Saline buffer (20 mM Tris, 137 mM NaCl, 0.1% Tween 20, pH 7.5) with 5% (w/v) skim milk powder as blocking reagent. SbtA protein was detected with a polyclonal antibody generated against a conserved 14 amino acid-epitope (Agrisera, Sweden). The HAHis-tagged SbtB proteins were detected with monoclonal antibodies against the HA-epitope (Sigma, USA). Secondary, alkaline phosphatase-conjugated antibodies (Bio-Rad, USA) were used for fluorometric detection of the immune-reactive bands with the AttoPhos system (Promega, USA). The fluorescence signal was visualized with a ChemiDoc™ MP imaging system (Biorad, USA). The relative signal intensities were quantified based on global background-corrected pixel volumes using ImageLab software (Bio-Rad, USA). Amounts of SbtA protein retained in complexes by binding to SbtB (pixel volumes; y) were plotted against adenylnucleotide ratios (AR; x), using OriginPro v2020 software (OriginLab Corp., USA). The non-linear correlation was fitted by a type 1 sigmoidal-logistic function (*F*(x) = y = A/(1 + e^-*k*\*(x-xc)^)), with A as maximum amount of SbtA bound*, k* as rate (binding) coefficient, and xc as the AR at the inflection point. The adenylnucleotide ratios corresponding to 50% (AR_50_) and 10% (AR_10_) of SbtA bound to SbtB were estimated from the sigmoidal logistic fitted curves normalized to residual SbtA detected at AR=1.

### Photosynthetic analysis by membrane inlet mass spectrometry (MIMS) measurements

Net photosynthetic O_2_ evolution and Ci uptake rates were calculated from changes in the concentrations of dissolved CO_2_ and O_2_ in aqueous cyanobacterial cell suspensions measured in a closed custom-built, temperature-controlled cuvette attached to a mass spectrometer (Isoprime IRMS series, Micromass, UK) as described previously (Badger and Andrews, 1982; Badger et al., 1985; Price et al., 2004). The inside of cuvette was separated from the inlet to the mass spectrometer by a gas permeable membrane. Low Ci-induced cell cultures were harvested by centrifugation (4 min, 4000 *g*, 23 °C), washed once with CO_2_-free BTP (1,3-bis[tris(hydroxymethyl)methylamino]propane) assay buffer (20 mM BTP buffered modified Bg-11(-NO_3_), 20 mM NaCl, pH 8.0), and resuspended at cell densities equal to 4 µg ml^-1^ Chl*a*. Chl*a* content was determined spectrophotometrically in 90% (v/v) methanol extracts (Porra et al., 1989). Measurements were performed at 30 °C which is equivalent to the growth temperature for cell cultures. During MIMS measurements, CO_2_ uptake was inhibited with 500 µM ethoxyzolamide (EZ). Steady state photosynthetic O_2_ evolution in response to light intensity was recorded for cell suspension cultures equilibrated with 1 mM HCO_3_^-^ which constitutes > 98% of the available Ci at pH 8. Illumination (KL1500 HAL light source, Schott, Germany) was increased stepwise from 0 to 15, 50, 200, 490, 850 and 1100 µmol photons m^-2^ s^-1^. Respiration rates were estimated from O_2_ consumption in the dark. Prior to MIMS measurements of Ci response curves, cell cultures were depleted for Ci by bubbling with N_2_ in white light (400 µmol m^-2^ s^-1^) for at least 10 min and photosynthetic O_2_ evolution rates recorded following incremental additions of bicarbonate. Steady state O_2_ evolution rates (*V*) normalized to Chl*a* contents were plotted against Ci substrate (S) concentrations, and resulting curves were fitted by the Michaelis-Menten equation, *V* = (*V*_max_ [S])/(*k*_0.5_ + [S], using OriginPro v2021 software. Maximum O_2_ evolution rates (*V*_maxO2_) were calculated as an approximation for maximum bicarbonate uptake by SbtA. Affinity of SbtA for HCO_3_ ^-^ was deduced from the Ci concentration supporting half-maximal activity (*k*_0.5Ci_). Effects of 300 s-illuminations with high (HL, 1400 µmol m^-2^ s^-1^) and low (LL, 200 µmol m^-2^ s^-1^) white light on HCO_3_ ^-^uptake and steady state O_2_ evolution rates were determined in cell cultures that had been equilibrated in the dark after addition of aqueous NaHCO_3_ solution to a final concentration of 250 or 500 µM.

### Ci uptake measurement by silicon oil centrifugation-filtration

Light- and dark stimulated uptake of active species of [^14^C]HCO_3_^-^ in cyanobacterial cell cultures was measured using the silicon oil centrifugation-filtration method as described previously (Kaplan et al., 1980; Price and Badger, 1989). Low Ci-induced cell cultures were prepared as described for MIMS analyses. Cells were pelleted by centrifugation (4 min, 4000 *g*, 23 °C) and resuspended in CO_2_-free BTP at pH 8 or pH 9 at densities equal to 4 µg ml^-1^ Chl*a*. Ci uptake was determined in 200 µl aliquots of cells supplemented with 500 µM EZ that were transferred into translucent centrifuge tubes on top of a silicon oil layer (mixture of AR200 : AR20 3.5 : 4 (v/v)) which separated the cells from the kill solution (50% methanol, 2 N NaOH) in the bottom of the tube. For low light (30 µmol photons m^-2^ s^-1^) activation experiments, cells were equilibrated with 500 µM [^14^C]HCO_3_^-^ substrate mix (aqueous NaHCO_3_ solution spiked with radioactive [^14^C]NaHCO_3_, pH 9.5) for 60 s either in the dark (dark control) or after exposure to low light for 30 or 60 s following a transfer from the dark. Uptake was terminated by rapid filtering of the cells into the kill solution during 20 s centrifugation at set maximum speed (microfuge B, Beckman, USA). For dark-inactivation experiments, cell cultures were depleted of Ci in the light (400 µmol m^-2^ s^-1^) prior to initiation of HCO_3_^-^ uptake by adding 500 µM [^14^C]HCO_3_^-^ substrate mix, either in the light (400 µmol m^-2^ s^-1^), or at various time points in the dark after cells had been transferred from the light into the dark for 15, 30, 45, 60, 90 and 180 s. Cells were incubated with ^14^C-labelled substrate for 30 s, and uptake was terminated by 20 s centrifugation. Tubes were immediately frozen in dry ice. The tips containing the cell pellets were cut off, resuspended in 2 N NaOH and mixed with 5 volumes scintillation fluid (Ultima Gold™ XR, PerkinElmer) for ^14^C-CPM counting in a TRI-CARB liquid scintillation counter (PerkinElmer, USA). Ci uptake rates were calculated based on the specific activity of the [^14^C]HCO_3_ ^-^ substrate mix. To test the inhibitory effect of EZ, cells were incubated with varying concentrations of EZ (0, 250, 375, 500, 750, 1000 µM) for 5 min in the light before adding 500 µM [^14^C]HCO_3_ ^-^ substrate mix for 30 s-uptake measurements.

### Bioinformatics

Homologous SbtA and SbtB proteins were identified in BLASTP (Altschul et al., 1990) searches and multiple pairwise amino acid sequence alignments generated in Geneious Prime 2021 (https://www.geneious.com) with MUSCLE software (Edgar, 2004). Further details on aligned proteins is given in **Supporting Information Tables S1** and **S2**. The three-dimensional domain homology models of the SbtB_7942_, SbtA_7942_ and SbtA_7001_ monomers were generated using AlphaFold2 (ColabFold v1.5.2: AlphaFold2 using MMseqs2) (Mirdita et al., 2022). Three-dimensional structure visualization and comparisons were performed with UCSF Chimera v.1.16 (http://www.cgl.ucsf.edu/chimera) (Meng et al., 2006) and UCSF ChimeraX (https://www.rbvi.ucsf.edu/chimerax) (Pettersen et al., 2021). The overall folds of SbtB_7942_ and SbtB_7001_ chain A with ADP (PDB: 6MMC) or Ca^2+^ and ATP ligands (PDB: 6MM2) proteins were compared by superposition onto each other and the cryo-EM structure of SbtB_6803_ (PDB: 7EGK, chains B, D, or F). The SbtA_7942_ and SbtA_7001_ homology models were compared by superposition onto each other and the cryo-EM structure of SbtA_6803_ (PDB: 7EGK, chains A, C or E).

### Statistics

Statistically significant differences between data sets were determined by performing a one-way analysis of variance (ANOVA) and post-hoc Tukey’s Honest Significant Difference (HSD) test (Tukey, 1949), using a type I error level α=0.05 (*P*=0.05) as threshold.

## Acknowledgements

This work was supported by the Australian Government through the Australian Research Council Centre of Excellence for Translational Photosynthesis (CE1401000015). Molecular graphics and analyses were performed with UCSF ChimeraX, developed by the Resource for Biocomputing, Visualization, and Informatics at the University of California, San Francisco, with support from National Institutes of Health R01-GM129325 and the Office of Cyber Infrastructure and Computational Biology, National Institute of Allergy and Infectious Diseases.

## Author Contributions

BF, BM and GDP: Conceptualization and Methodology. BF, BM and LR: Investigation. BF: Formal data analysis. BF: Writing - original manuscript draft. All authors: Writing-review & editing. GDP: Funding acquisition.

## Data availability

All data supporting this study are available within the main manuscript and the supplementary information published online. Any data not shown are available from the corresponding author upon request.

## References

Agostoni M, Montgomery BL (2014) Survival strategies in the aquatic and terrestrial world: The impact of second messengers on cyanobacterial processes. Life 4: 745–769

Altschul SF, Gish W, Miller W, Myers EW, Lipman DJ (1990) Basic local alignment search tool. Journal of Molecular Biology 215: 403–410

Angeleri M, Muth-Pawlak D, Aro E-M, Battchikova N (2016) Study of O-phosphorylation sites in proteins involved in photosynthesis-related processes in *Synechocystis sp.* strain PCC 6803: application of the SRM approach. Journal of Proteome Research 15: 4638–4652

Atkinson DE, Walton GM (1967) Adenosine triphosphate conservation in metabolic regulation: Rat liver citrate cleavage enzyme. Journal of Biological Chemistry 242: 3239–3241

Badger MR, Andrews TJ (1982) Photosynthesis and inorganic carbon usage by the marine cyanobacterium, *Synechococcus sp*. Plant Physiology 70: 517–523

Badger MR, Bassett M, Comins HN (1985) A model for HCO_3_^-^ accumulation and photosynthesis in the cyanobacterium *Synechococcus sp*: Theoretical predictions and experimental observations. Plant Physiology 77: 465–471

Badger MR, Price GD (1992) The CO_2_ concentrating mechanism in cyanobactiria and microalgae. Physiologia Plantarum 84: 606–615

Bornefeld T, Simonis W (1974) Effects of light, temperature, pH, and inhibitors on the ATP level of the blue-green alga *Anacystic nidulans*. Planta 115: 309–318

Cann M (2004) Bicarbonate stimulated adenylyl cyclases. IUBMB Life 56: 529–534

Chapman AG, Fall L, Atkinson DE (1971) Adenylate energy charge in *Escherichia coli* during growth and starvation. Journal of Bacteriology 108: 1072–1086

De la Fuente IM, Cortés JM, Valero E, Desroches M, Rodrigues S, Malaina I, Martínez L (2014) On the dynamics of the adenylate energy system: homeorhesis vs homeostasis. PLOS ONE 9: e108676

Du J, Förster B, Rourke L, Howitt SM, Price GD (2014) Characterisation of cyanobacterial bicarbonate transporters in *E. Coli* shows that SbtA homologs are functional in this heterologous expression system. PLoS ONE 9: e115905

Dzelzkalns VA, Owens GC, Bogorad L (1984) Chloroplast promoter driven expression of the chloramphenicol acetyl transferase gene in a cyanobacterium. Nucleic acids research 12: 8917–8925

Edgar RC (2004) MUSCLE: a multiple sequence alignment method with reduced time and space complexity. BMC Bioinformatics 5: 113

Espie G, Kimber M (2011) Carboxysomes: cyanobacterial RubisCO comes in small packages. Photosynthesis Research 109: 7–20

Fang S, Huang X, Zhang X, Zhang M, Hao Y, Guo H, Liu L-N, Yu F, Zhang P (2021) Molecular mechanism underlying transport and allosteric inhibition of bicarbonate transporter SbtA. Proceedings of the National Academy of Sciences 118: e2101632118

Field CB, Behrenfeld MJ, Randerson JT, Falkowski P (1998) Primary production of the biosphere: Integrating terrestrial and oceanic components. Science 281: 237–240

Forchhammer K, Lüddecke J (2016) Sensory properties of the PII signalling protein family. The FEBS Journal 283: 425–437

Forchhammer K, Selim KA (2020) Carbon/nitrogen homeostasis control in cyanobacteria. FEMS Microbiology Reviews 44: 33–53

Golden SS, Sherman LA (1984) Optimal conditions for genetic transformation of the cyanobacterium Anacystis nidulans R2. Journal of Bacteriology 158: 36–42

Hammer A, Hodgson DRW, Cann MJ (2006) Regulation of prokaryotic adenylyl cyclases by CO2. The Biochemical Journal 396: 215–218

Huergo LF, Chandra G, Merrick M (2013) PII signal transduction proteins: nitrogen regulation and beyond. FEMS Microbiology Reviews 37: 251–283

Huergo LF, Dixon R (2015) The emergence of 2-oxoglutarate as a master regulator metabolite. Microbiology and Molecular Biology Reviews 79: 419–435

Kaczmarski JA, Hong N-S, Mukherjee B, Wey LT, Rourke L, Förster B, Peat TS, Price GD, Jackson CJ (2019) Structural basis for the allosteric regulation of the SbtA bicarbonate transporter by the PII-like protein, SbtB, from Cyanobium sp. PCC7001. Biochemistry 58: 5030-5039

Kallas T, Castenholz RW (1982) Internal pH and ATP-ADP pools in the cyanobacterium *Synechococcus sp.* during exposure to growth-inhibiting low pH. Journal of Bacteriology 149: 229–236

Kaplan A, Badger M, Berry J (1980) Photosynthesis and the intracellular inorganic carbon pool in the bluegreen alga *Anabaena variabilis*: Response to external CO_2_ concentration. Planta 149: 219–226

Kaplan A, Reinhold L (1999) CO_2_ concentrating mechanisms in photosynthetic microorganisms. Annual Review of Plant Physiology and Plant Molecular Biology 50: 539–570

Kaplan A, Zenvirth D, Marcus Y, Omata T, Ogawa T (1987) Energization and activation of inorganic carbon uptake by light in cyanobacteria. Plant Physiology 84: 210–213

Leganés F, Forchhammer K, Fernández-Piñas F (2009) Role of calcium in acclimation of the cyanobacterium *Synechococcus elongatus* PCC 7942 to nitrogen starvation. Microbiology 155: 25–34

Liu H, Nolla H, Campbell L (1997) Prochlorococcus growth rate and contribution to primary production in the equatorial and subtropical North Pacific Ocean. Aquatic Microbial Ecology 12: 39–47

Liu X-Y, Hou W-T, Wang L, Li B, Chen Y, Chen Y, Jiang Y-L, Zhou C-Z (2021) Structures of cyanobacterial bicarbonate transporter SbtA and its complex with PII-like SbtB. Cell Discovery 7: 63

Long BM, Rae BD, Rolland V, Förster B, Price GD (2016) Cyanobacterial CO_2_-concentrating mechanism components: function and prospects for plant metabolic engineering. Current Opinion in Plant Biology 31: 1–8

Maeda S-i, Badger MR, Price GD (2002) Novel gene products associated with NdhD3/D4-containing NDH-1 complexes are involved in photosynthetic CO2 hydration in the cyanobacterium, Synechococcus sp. PCC7942. Molecular Microbiology 43: 425-435

McGinn PJ, Price GD, Maleszka R, Badger MR (2003) Inorganic carbon limitation and light control the expression of transcripts related to the CO_2_-concentrating mechanism in the cyanobacterium *Synechocystis sp.* strain PCC6803. Plant Physiology 132: 218–229

McGrath JM, Long SP (2014) Can the cyanobacterial carbon-concentrating mechanism increase photosynthesis in crop species? A theoretical analysis. Plant Physiology 164: 2247–2261

Meng EC, Pettersen EF, Couch GS, Huang CC, Ferrin TE (2006) Tools for integrated sequence-structure analysis with UCSF Chimera. BMC Bioinformatics 7: 339–339

Meyer MT, McCormick AJ, Griffiths H (2016) Will an algal CO_2_-concentrating mechanism work in higher plants? Current Opinion in Plant Biology 31: 181–188

Mirdita M, Schütze K, Moriwaki Y, Heo L, Ovchinnikov S, Steinegger M (2022) ColabFold: making protein folding accessible to all. Nature Methods 19: 679–682

Ogawa T, Miyano A, Inoue Y (1985) Photosystem-I-driven inorganic carbon transport in the cyanobacterium, Anacystis nidulans. Biochimica et Biophysica Acta (BBA) - Bioenergetics 808: 77–84

Ohkawa H, Sonoda M, Hagino N, Shibata M, Pakrasi HB, Ogawa T (2002) Functionally distinct NAD(P)H dehydrogenases and their membrane localization in Synechocystis sp. PCC6803. Functional Plant Biology 29: 195-200

Ohmori M, Okamoto S (2004) Photoresponsive cAMP signal transduction in cyanobacteria. Photochemical & Photobiological Sciences 3: 503–511

Omata T, Carlson TJ, Ogawa T, Pierce J (1990) Sequencing and Modification of the Gene Encoding the 42-Kilodalton Protein in the Cytoplasmic Membrane of Synechococcus PCC 7942. Plant physiology 93: 305–311

Omata T, Price GD, Badger MR, Okamura M, Gohta S, Ogawa T (1999) Identification of an ATP-binding cassette transporter involved in bicarbonate uptake in the cyanobacterium *Synechococcus sp.* strain PCC 7942. Proceedings of the National Academy of Sciences 96: 13571–13576

Oren N, Timm S, Frank M, Mantovani O, Murik O, Hagemann M (2021) Red/far-red light signals regulate the activity of the carbon-concentrating mechanism in cyanobacteria. Science Advances 7: eabg0435

Pettersen EF, Goddard TD, Huang CC, Meng EC, Couch GS, Croll TI, Morris JH, Ferrin TE (2021) UCSF ChimeraX: Structure visualization for researchers, educators, and developers. Protein Sci 30: 70–82

Porra RJ, Thompson WA, Kriedemann PE (1989) Determination of accurate extinction coefficients and simultaneous equations for assaying chlorophylls a and b extracted with four different solvents: verification of the concentration of chlorophyll standards by atomic absorption spectroscopy. Biochimica et Biophysica Acta (BBA) - Bioenergetics 975: 384–394

Price G (2011) Inorganic carbon transporters of the cyanobacterial CO_2_ concentrating mechanism. Photosynthesis Research 109: 47–57

Price GD, Badger MR (1989) Ethoxyzolamide inhibition of CO_2_ uptake in the cyanobacterium *Synechococcus* PCC7942 without apparent inhibition of internal carbonic anhydrase activity. Plant Physiology 89: 37–43

Price GD, Badger MR (1989) Isolation and characterization of high CO_2_-requiring-mutants of the cyanobacterium *Synechococcus* PCC7942: Two phenotypes that accumulate inorganic carbon but are apparently unable to generate CO_2_ within the carboxysome. Plant Physiology 91: 514–525

Price GD, Badger MR, Woodger FJ, Long BM (2008) Advances in understanding the cyanobacterial CO_2_-concentrating-mechanism (CCM): functional components, Ci transporters, diversity, genetic regulation and prospects for engineering into plants. Journal of Experimental Botany 59: 1441–1461

Price GD, Pengelly JJL, Förster B, Du J, Whitney SM, von Caemmerer S, Badger MR, Howitt SM, Evans JR (2013) The cyanobacterial CCM as a source of genes for improving photosynthetic CO_2_ fixation in crop species. Journal of Experimental Botany 64: 753–768

Price GD, Shelden MC, Howitt SM (2011) Membrane topology of the cyanobacterial bicarbonate transporter, SbtA, and identification of potential regulatory loops. Molecular Membrane Biology 28: 265–275

Price GD, Woodger FJ, Badger MR, Howitt SM, Tucker L (2004) Identification of a SulP-type bicarbonate transporter in marine cyanobacteria. Proceedings of the National Academy of Sciences 101: 18228–18233

Radchenko MV, Thornton J, Merrick M (2010) Control of AmtB-GlnK complex formation by intracellular levels of ATP, ADP, and 2-oxoglutarate. Journal of Biological Chemistry 285: 31037-31045

Radchenko MV, Thornton J, Merrick M (2013) P_II_ signal transduction proteins are ATPases whose activity is regulated by 2-oxoglutarate. Proceedings of the National Academy of Sciences 110: 12948–12953

Rae BD, Förster B, Badger MR, Price GD (2011) The CO_2_-concentrating mechanism of *Synechococcus* WH5701 is composed of native and horizontally-acquired components. Photosynthesis Research 109: 59–72

Rath A, Glibowicka M, Nadeau VG, Chen G, Deber CM (2009) Detergent binding explains anomalous SDS-PAGE migration of membrane proteins. Proceedings of the National Academy of Sciences 106: 1760–1765

Rust MJ, Golden SS, O’Shea EK (2011) Light-driven changes in energy metabolism directly entrain the cyanobacterial circadian oscillator. Science 331: 220–223

Sánchez-Baracaldo P, Bianchini G, Di Cesare A, Callieri C, Chrismas NAM (2019) Insights into the evolution of picocyanobacteria and phycoerythrin genes (*mpeBA* and *cpeBA*). Frontiers in Microbiology 10: 45

Schwarz D, Nodop A, Hüge J, Purfürst S, Forchhammer K, Michel K-P, Bauwe H, Kopka J, Hagemann M (2011) Metabolic and transcriptomic phenotyping of inorganic carbon acclimation in the cyanobacterium *Synechococcus elongatus* PCC 7942. Plant Physiology 155: 1640–1655

Selim KA, Haase F, Hartmann MD, Hagemann M, Forchhammer K (2018) P_II_-like signaling protein SbtB links cAMP sensing with cyanobacterial inorganic carbon response. Proceedings of the National Academy of Sciences 115: E4861–E4869

Selim KA, Haffner M, Burkhardt M, Mantovani O, Neumann N, Albrecht R, Seifert R, Krüger L, Stülke J, Hartmann MD, Hagemann M, Forchhammer K (2021) Diurnal metabolic control in cyanobacteria requires perception of second messenger signaling molecule c-di-AMP by the carbon control protein SbtB. Science Advances 7: eabk0568

Selim KA, Haffner M, Mantovani O, Albrecht R, Zhu H, Hagemann M, Forchhammer K, Hartmann MD (2023) Carbon signaling protein SbtB possesses atypical redox-regulated apyrase activity to facilitate regulation of bicarbonate transporter SbtA. Proceedings of the National Academy of Sciences 120: e2205882120

Selim KA, Haffner M, Watzer B, Forchhammer K (2019) Tuning the in vitro sensing and signaling properties of cyanobacterial PII protein by mutation of key residues. Scientific Reports 9: 18985

Shibata M, Katoh H, Sonoda M, Ohkawa H, Shimoyama M, Fukuzawa H, Kaplan A, Ogawa T (2002) Genes essential to sodium-dependent bicarbonate transport in cyanobacteria. Journal of Biological Chemistry 277: 18658–18664

Spät P, Maček B, Forchhammer K (2015) Phosphoproteome of the cyanobacterium Synechocystis sp. PCC 6803 and its dynamics during nitrogen starvation. Frontiers in Microbiology 6

Sültemeyer D, Klughammer B, Badger MR, Dean Price G (1998) Fast induction of high-affinity HCO_3_ ^−^ transport in cyanobacteria. Plant Physiology 116: 183–192

Takano S, Tomita J, Sonoike K, Iwasaki H (2015) The initiation of nocturnal dormancy in Synechococcus as an active process. BMC Biology 13: 36

Tamoi M, Miyazaki T, Fukamizo T, Shigeoka S (2005) The Calvin cycle in cyanobacteria is regulated by CP12 via the NAD(H)/NADP(H) ratio under light/dark conditions. The Plant Journal 42: 504–513

Thauer RK, Jungermann K, Decker K (1977) Energy conservation in chemotrophic anaerobic bacteria. Bacteriological Reviews 41: 100–180

Torrecilla I, Leganés F, Bonilla I, Fernández-Piñas F (2004) Light-to-dark transitions trigger a transient increase in intracellular Ca2+ modulated by the redox state of the photosynthetic electron transport chain in the cyanobacterium Anabaena sp. PCC7120. Plant, Cell & Environment 27: 810-819

Tukey JW (1949) Comparing individual means in the analysis of variance. Biometrics 5: 99–114

Wheatley NM, Eden KD, Ngo J, Rosinski JS, Sawaya MR, Cascio D, Collazo M, Hoveida H, Hubbell WL, Yeates TO (2016) A PII-like protein regulated by bicarbonate: Structural and biochemical studies of the carboxysome-associated CPII protein. Journal of Molecular Biology 428: 4013–4030

Woodger FJ, Badger MR, Price GD (2003) Inorganic carbon limitation induces transcripts encoding components of the CO2-concentrating mechanism in Synechococcus sp. PCC7942 through a redox-independent pathway. Plant Physiology 133: 2069-2080

Xu M, Su Z (2009) Computational prediction of cAMP receptor protein (CRP) binding sites in cyanobacterial genomes. BMC Genomics 10: 23

Yaginuma H, Kawai S, Tabata KV, Tomiyama K, Kakizuka A, Komatsuzaki T, Noji H, Imamura H (2014) Diversity in ATP concentrations in a single bacterial cell population revealed by quantitative single-cell imaging. Scientific reports 4: 6522–6522

Zeth K, Fokina O, Forchhammer K (2014) Structural basis and target-specific modulation of ADP sensing by the *Synechococcus elongatus* PII signaling protein. The Journal of Biological Chemistry 289: 8960–8972

