## Supplemental information for "Adenylnucleotide-mediated binding of the PII-like protein SbtB contributes to controlling activity of the cyanobacterial bicarbonate transporter SbtA"

**For manuscript:**

**Fig. S1** Membrane-intrinsic SbtA and soluble SbtB proteins both accumulate in membrane enriched fractions.

**Fig. S2** The association of SbtA and cognate SbtB proteins is not facilitated by $Mg^{2+}$ or $Ca^{2+}$ ions and in combination with $HCO_3^-$.

**Fig. S3** Bicarbonate levels do not affect the association of SbtA and SbtB proteins.

**Fig. S4** Quantification of SbtA:SbtB binding in response to adenylnucleotide ratios simulating cellular adenylate energy charge variations.

**Fig. S5** SbtA-mediated bicarbonate uptake in *Synechococcus elongatus* PCC7942 is not affected by the carbonic anhydrase inhibitor ethoxyzolamide.

**Fig. S6** Net photosynthetic $O_2$ evolution rates of *Synechococcus* ΔCS supported by SbtA-dependent bicarbonate uptake in the presence and absence of SbtB.

**Fig. S7** Effects of changes in light intensity on SbtA-dependent photosynthesis rates.

**Fig. S8** Growth of *Synechococcus* ΔCS cells expressing $SbtA_{7942}$ (SbtA) compared to cells co-expressing $SbtA_{7942}$ with SbtB7942 (SbtAB) or with HAHis-tagged SbtB7942 (SbtABH).

**Fig. S9** Amino acid sequence alignment of the SbtA proteins and SbtB proteins from *Cyanobium sp.* PCC7001, *Synechococcus elongatus* PCC7942 and *Synechocystis sp.* PCC6803.

**Fig. S10** Amino acid alignment of SbtA proteins homologs ordered by similarity to $SbtA_{7001}$.

**Fig. S11** Amino acid alignment of SbtB protein homologs ordered by similarity to $SbtB_{7001}$.

**Fig. S12** Structural comparison of the overall fold of SbtA and SbtB proteins from *Synechococcus elongatus* PCC7942, *Cyanobium sp.* PCC7001 and *Synechocystis sp.* PCC6803.

**Table S1** List of gene identifiers for SbtA protein homologs aligned in Fig. S10.

**Table S2** List of gene identifiers for SbtB protein homologs aligned in Fig. S11.

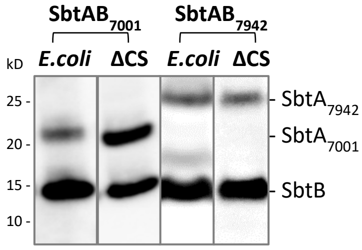


**Fig. S1. Membrane-intrinsic SbtA and soluble SbtB proteins both accumulate in membrane-enriched fractions.** The SbtA-SbtB pairs of *Cyanobium sp.* PCC7001 ($SbtA_{7001}$ and $SbtB_{7001}$, respectively) and *Synechococcus elongatus* PCC7942 ($SbtA_{7942}$ and $SbtB_{7942}$, respectively) were expressed from plasmids in *E. coli* and in *Synechococcus* ΔCS. The membrane-enriched, native protein extracts, solubilized in 1% DDM, represent typical preparations used in IMAC binding assays. Immunological detection of the HAHis-tagged SbtB proteins and SbtA proteins showed substantial amounts of both the membrane-integral SbtA protein and the cytoplasmic SbtB proteins were present in the membrane-enriched fraction. This was routinely tested to ensure protein preparations were suitable to investigate the physical interaction between SbtA and SbtB. Note that on SDS-PAGE, apparent and calculated molecular weights match for both HAHis-tagged SbtB monomers (13 kD), whereas the apparent molecular weights of both $SbtA_{7001}$ and $SbtA_{7942}$ proteins are consistently lower than their calculated molecular weights (23 vs 35 kD and 29 vs 40 kD, respectively). Representative results are shown as composite images from different Western blots.

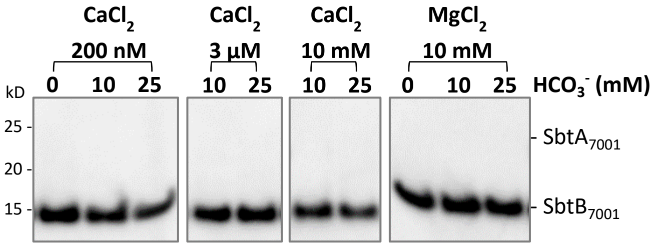


**Fig. S2. The association of SbtA and SbtB proteins is not facilitated by $Ca^{2+}$ or $Mg^{2+}$ ions and in combination with $HCO_3^-$.** The $SbtA_{7001}$-$SbtB_{7001}$ protein pair was expressed in *E. coli*. In IMAC binding assays, membrane-enriched native protein extracts were incubated with 200 nM, 3 µM, 10 mM $CaCl_2$ or 10 mM $MgCl_2$ and combined with 10 or 25 mM $HCO_3^-$. The binding assay recovered SbtBHAHis irrespective of the effectors added as it binds directly to the beads, whereas SbtA does not. Therefore, coelution of SbtA was evidence for its interaction with SbtBHAHis. Immunological detection of the HAHis-tagged SbtB proteins and SbtA proteins showed SbtA:SbtB association was not promoted by adding $Mg^{2+}$, $Ca^{2+}$ and $Cl^-$ ions. Similarly, the presence of $HCO_3^-$ as the substrate for the SbtA transporter did not affect the SbtA:SbtB association. Thus, it could be ruled out that the 200 nM $CaCl_2$ that were added to IMAC binding assays influenced SbtA:SbtB binding.

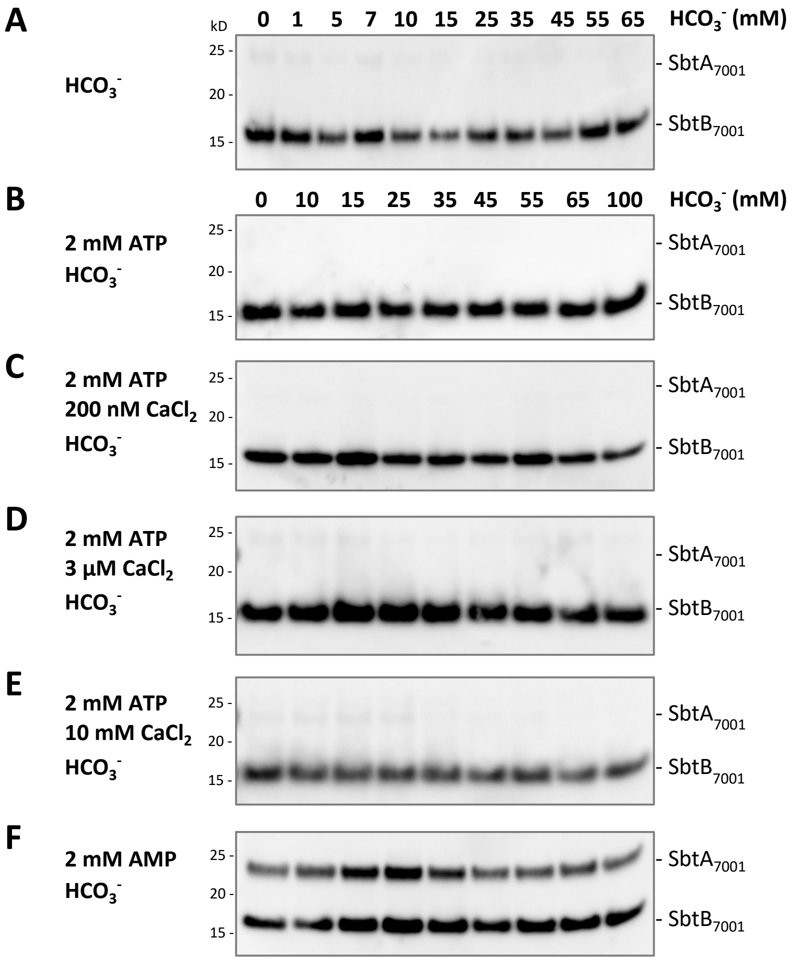


**Fig. S3. Bicarbonate levels do not affect the association of SbtA and SbtB proteins.** The $SbtA_{7001}$-$SbtB_{7001}$ protein pair was expressed in *E. coli*. In IMAC binding assays, membrane-enriched native protein extracts were incubated with (**A**) no $CaCl_2$ and 0 to 100 mM $NaHCO_3$ ($HCO_3^-$), (**B**) no $CaCl_2$ and 2 mM ATP, (**C**) 200 nM $CaCl_2$ and 2 mM ATP, (**D**) 3 µM $CaCl_2$ and 2 mM ATP, (**E**) 10 mM $CaCl_2$ and 2 mM ATP, (**F**) 2 mM AMP. The SbtBHAHis and SbtA proteins were detected immunologically on Western blots. Physiological levels (and above) of the substrate, $HCO_3^-$, did not alter SbtA:SbtB binding by itself and in combination with other effectors, ruling out synergistic effects between $HCO_3^-$ and the other effectors tested. SbtA remained dissociated from SbtB in the presence of ATP, irrespective of the presence of $Ca^{2+}$even though $Ca^{2+}$ was shown to enhance the stabilizing effect of ATP on the T-loop of $SbtB_{7001}$ and T-loop mediated protein-protein interaction. Similarly, SbtA and SbtB formed a complex in the presence of AMP independent of $HCO_3^-$.

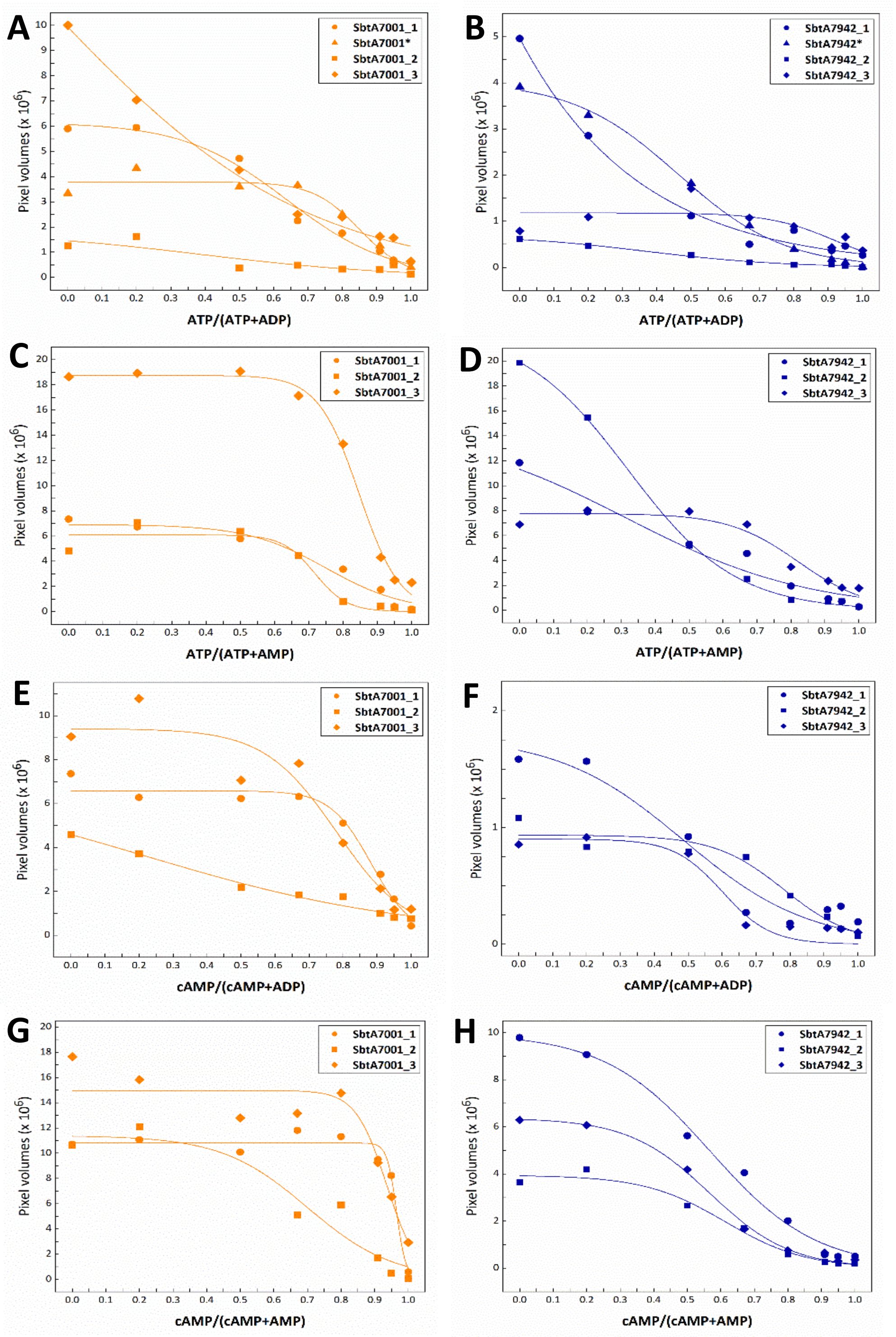


**Fig. S4. Quantification of SbtA:SbtB binding in response to adenylnucleotide ratios that simulate cellular adenylate energy charge variations.** Amounts of SbtA bound to SbtB protein at different ATP:ADP (**A**, **B**), ATP:AMP (**C**, **D**), cAMP:ADP (**E**, **F**) and cAMP:AMP (**G**, **H**) ratios were determined from *in vitro* IMAC binding assays. The SbtA$_{7001}$-SbtB$_{7001}$ (**A**, **C**, **E**, **G**) and SbtA$_{7942}$-SbtB$_{7942}$ (**B**, **D**, **F**, **H**) pairs were expressed in *E. coli* and *Synechococcus* ΔCS. The amount of SbtA within the SbtA:SbtB complexes was quantified by densitometry of Western blots images (pixel volumes) and plotted against the adenylnucleotide ratio. The correlation was non-linear and fitted by a sigmoidal-logistic function. The adjusted $R^2$ statistics indicated the fit quality (better fits nearing a value of 1). (**A**) SbtA7001_1, *, 2, 3 ($R^2$= 0.982, 0.950, 0.689, 0.986); (**B**) SbtA7942_1, *, 2, 3 ($R^2$= 0.988, 0.996, 0.987, 0.440); (**C**) SbtA7001_1, 2, 3 ($R^2$= 0.958, 0.933, 0.993); (**D**) SbtA7942_1, 2, 3 ($R^2$= 0.942, 0.999, 0.937); (**E**) SbtA7001_1, 2, 3 ($R^2$= 0.968, 0.973, 0.914); (**F**) SbtA7942_1, 2, 3 ($R^2$= 0.928, 0.939, 0.908); (**G**) SbtA7001_1, 2, 3 ($R^2$= 0.953, 0.890, 0.871); (**H**) SbtA7942_1, 2, 3 ($R^2$= 0.988, 0.979, 0.994); average adjusted $R^2$ ± SE = 0.926 ± 0.023.

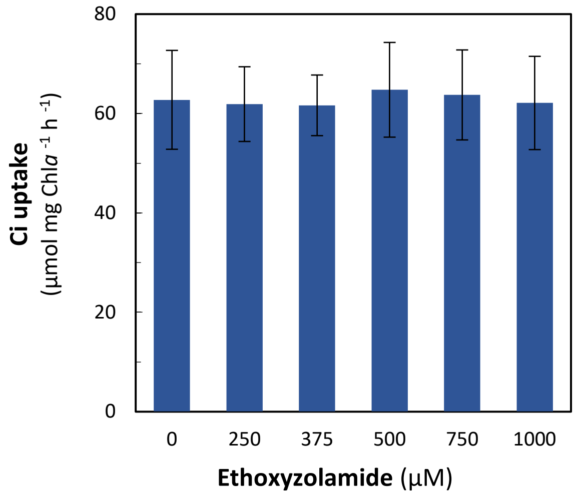


**Fig. S5. SbtA-mediated bicarbonate uptake in *Synechococcus elongatus* PCC7942 is not affected by the carbonic anhydrase inhibitor ethoxyzolamide (EZ).** As a control experiment, effects of EZ on total bicarbonate uptake by $SbtA_{7942}$, which is the intracellular accumulation of photosynthetically fixed $^{14}C$ and a free $[^{14}C]HCO_3^-$ pool, were measured in the $HCO_3^-$ transporter deletion strain *Synechococcus* ΔCS expressing $SbtA_{7942}$ from a plasmid using the silicon oil centrifugation-filtration method. Ci-depleted cell cultures, supplemented with various concentrations of EZ, were incubated in 500 µM $[^{14}C]HCO_3^-$ substrate mix at pH 8 for 30 s in the light (400 µmol photons $m^{-2}$ $s^{-1}$). The $SbtA_{7942}$-specific $HCO_3^-$ uptake rates were obtained by subtraction of the ΔCS background Ci uptake rates which represent the unspecific incorporation of $[^{14}C]$ label from the SbtA Ci uptake rate. Unspecific background label is mainly due to extracellular $[^{14}C]HCO_3^-$ trapped in the water shell surrounding each cell and a very small amount of $^{14}CO_2$ diffusion into the cells. Values are means ± SE from three independent experiments. Importantly, bicarbonate uptake by $SbtA_{7942}$ was unchanged across the concentration range of EZ tested, which ruled out any variations in bicarbonate uptake activity were caused by EZ that was routinely added to specifically inhibit the $CO_2$ uptake system during functional measurements.

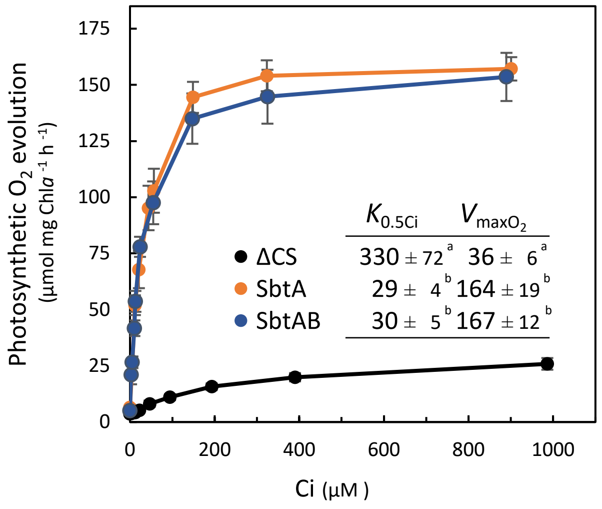


**Fig. S6. Net photosynthetic $O_2$ evolution rates of *Synechococcus* ΔCS supported by SbtA-dependent bicarbonate uptake in the presence and absence of SbtB.** The $SbtA_{7942}$ protein was expressed without (SbtA) or with $SbtB_{7942}$ (SbtAB) protein from plasmids in low Ci-induced *Synechococcus* ΔCS. Steady state photosynthetic $O_2$ evolution rates were determined over a range of external Ci concentrations in white light (400 µmol photons $m^{-2}$ $s^{-1}$), while $CO_2$ uptake was inhibited by EZ, using MIMS. Ci was supplied as $HCO_3^-$ which was added in increments to $CO_2$-depleted cell suspensions. Maximum $O_2$ evolution rates ($V_{maxO_2}$, µmol mg Chl*a*$^{-1}$ $h^{-1}$) and the Ci concentrations required for half-maximum photosynthetic rates ($K_{0.5Ci}$, µM) were derived from Michaelis-Menten kinetics (inset). Note that $HCO_3^-$ contributed 98% of the total Ci pool which means the actual substrate affinities for SbtA and SbtAB were 28 ± 4 and 29 ± 5, respectively. Statistically significant differences ($P$<0.05) are indicated by different letters. Values are means ± SE; n=11 (ΔCS), n=9 (SbtA), n=10 (SbtAB). The presence or absence of SbtB does not affect light-saturated photosynthetic rates that are almost entirely determined by SbtA-mediated $HCO_3^-$ uptake under these experimental conditions.

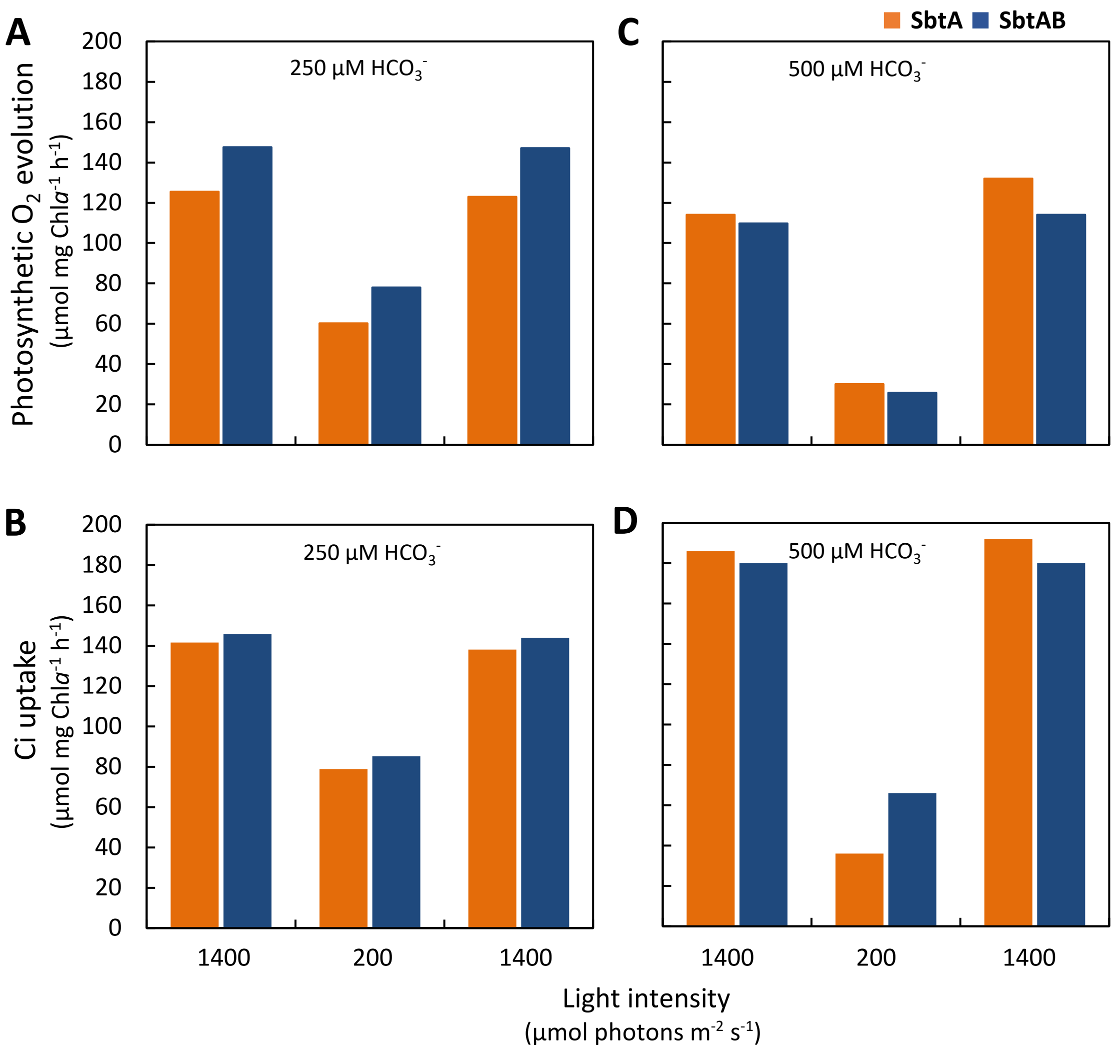


**Fig. S7**. **Effects of changes in light intensity on SbtA-dependent photosynthesis rates.**
Photosynthetic $O_2$ evolution (**A**, **C**) and $HCO_3^-$ uptake (**B**, **D**) rates of low Ci-induced *Synechococcus* ΔCS expressing $SbtA_{7942}$ only (SbtA), or co-expressing $SbtA_{7942}$ and $SbtB_{7942}$ (SbtAB) from plasmids. Photosynthetic $O_2$ evolution and Ci uptake rates were determined at 250 (**A**, **B**) and 500 µM (**C**, **D**) $HCO_3^-$ added to Ci-depleted cell suspensions in the dark before exposure to light, using MIMS. To ensure $HCO_3^-$ was the only Ci species effectively taken up by the cells, measurements were conducted at pH 8 and $CO_2$ uptake was inhibited adding 500 µM EZ. Therefore, Ci uptake was directly proportional to $HCO_3^-$ uptake by SbtA. Cells were exposed to 1400 µmol $m^{-2}$ $s^{-1}$ white light for about 5 min until Ci uptake and $O_2$ evolution rates were steady, followed by 5 min illumination with 200 µmol $m^{-2}$ $s^{-1}$ and again 5 min illumination with 1400 µmol $m^{-2}$ $s^{-1}$. Steady state $O_2$ evolution and Ci uptake rates normalized to Chl*a* were very similar between SbtA and SbtAB, irrespective of light intensity and Ci supplied, suggesting $SbtB_{7942}$ was likely not required for short term adjustments of photosynthesis and $SbtA_{7942}$ activity to light intensity and Ci availability.

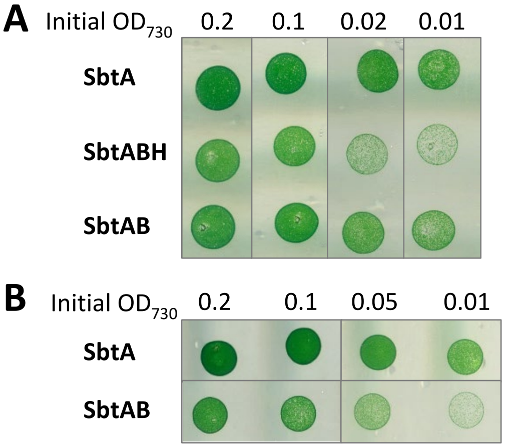


**Fig. S8. Growth of *Synechococcus* ΔCS cells expressing $SbtA_{7942}$ (SbtA) compared to cells co-expressing $SbtA_{7942}$ with $SbtB_{7942}$ (SbtAB) or with HAHis-tagged $SbtB_{7942}$ (SbtABH).** SbtA and SbtB proteins were expressed from plasmid-located genes under control of their native promoters in cell suspensions bubbled with air (ambient $CO_2$) for 4 h which fully induced the CCM. Cells suspensions adjusted to $OD_{730}$ of 0.2 and 1:2, 1:10 and 1:20 dilutions were spotted onto modified BG-11 agar plates (pH 8) for growth in air under diurnal (12 h light:12 h dark) cycles. Representative images of cell patches on day 10 from two separate experiments, (**A**) and (**B**), are shown. The composite images consists of selected areas from the same growth plate for each experiment. Cell density was evaluated visually, and indicated that cells accumulated the same or slightly more biomass when $SbtA_{7942}$ was expressed without the $SbtB_{7942}$ protein in low Ci-acclimated cells. This suggests SbtB was not required for optimal growth of *Synechococcus ΔCS* in low Ci when Ci was provided by both the $CO_2$ pumps $NDH1_{3/4}$ and bicarbonate uptake by SbtA.

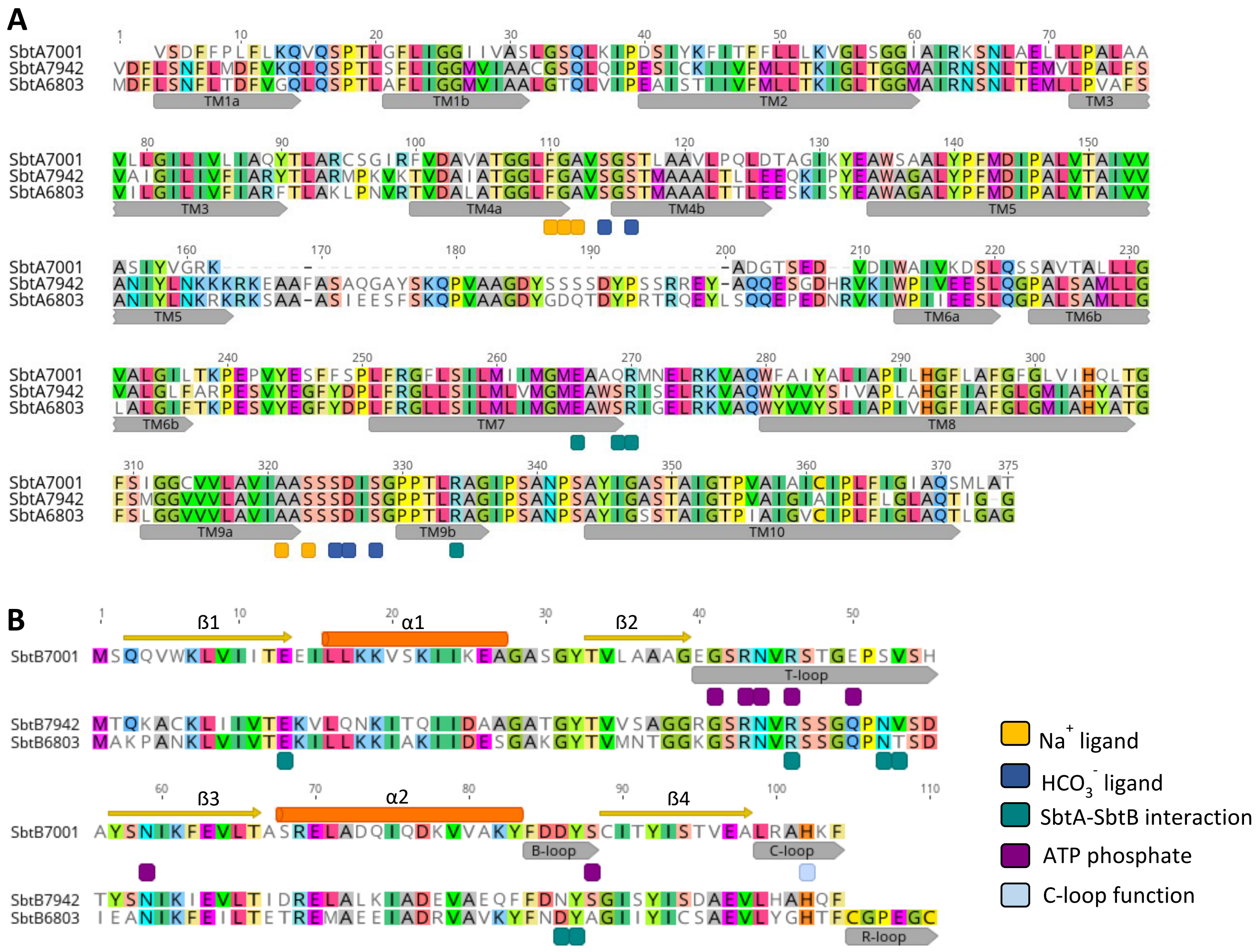


**Fig. S9. Amino acid sequence alignment of the SbtA proteins and SbtB proteins from *Cyanobium sp.* PCC7001, *Synechococcus elongatus* PCC7942 and *Synechocystis sp.* PCC6803.** Identical residues are highlighted in colour. (**A**) Alignment of SbtA proteins. Transmembrane domains (TM; grey arrows), amino acids coordinating $Na^+$ and $HCO_3^-$ and residues implicated in SbtA-SbtB interaction are annotated based on sequence homology and predictions from cryo-EM structures of $SbtA_{6803}$ (PDB: 7EGK). The $SbtA_{7001}$ and $SbtA_{7942}$ proteins share 60% overall identity. (**B**) Alignment of SbtB proteins. The T-, B-, C-, and R-loops (grey arrows) and amino acids important for interaction with ATP phosphates and conformation of T- and C-loops are based on the crystal structure of $SbtB_{7001}$ (PDB: 6MM2) and cryo-EM structure of $SbtB_{6803}$ (PDB: 7EGK). Similarly, the $SbtB_{7001}$ and $SbtB_{7942}$ share 57.7% overall identity.

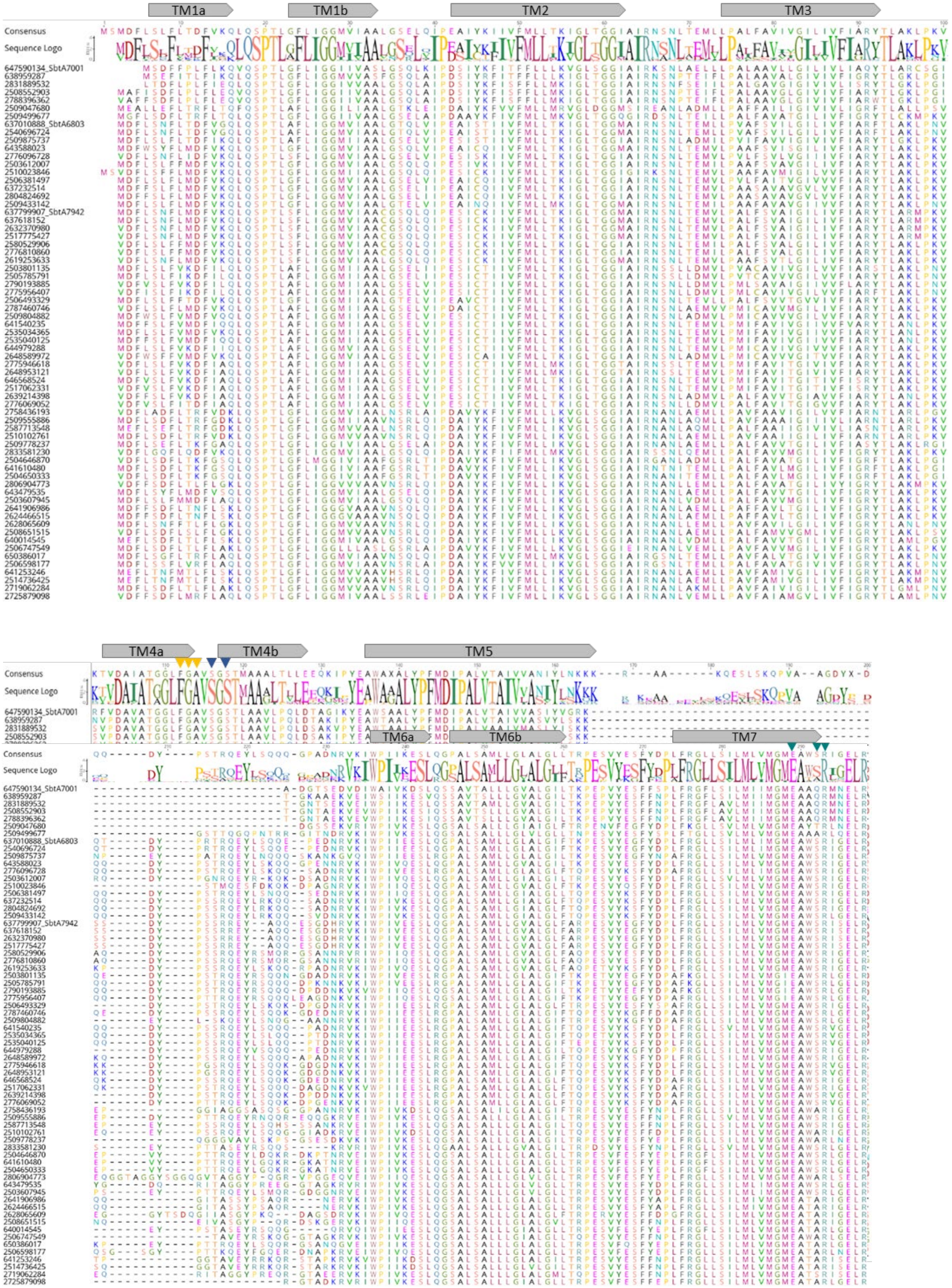
TM1a
TM1b
TM2
TM3
Consensus
Sequence Logo
TM4a
TM4b
TM5
TM6a
TM6b
TM7

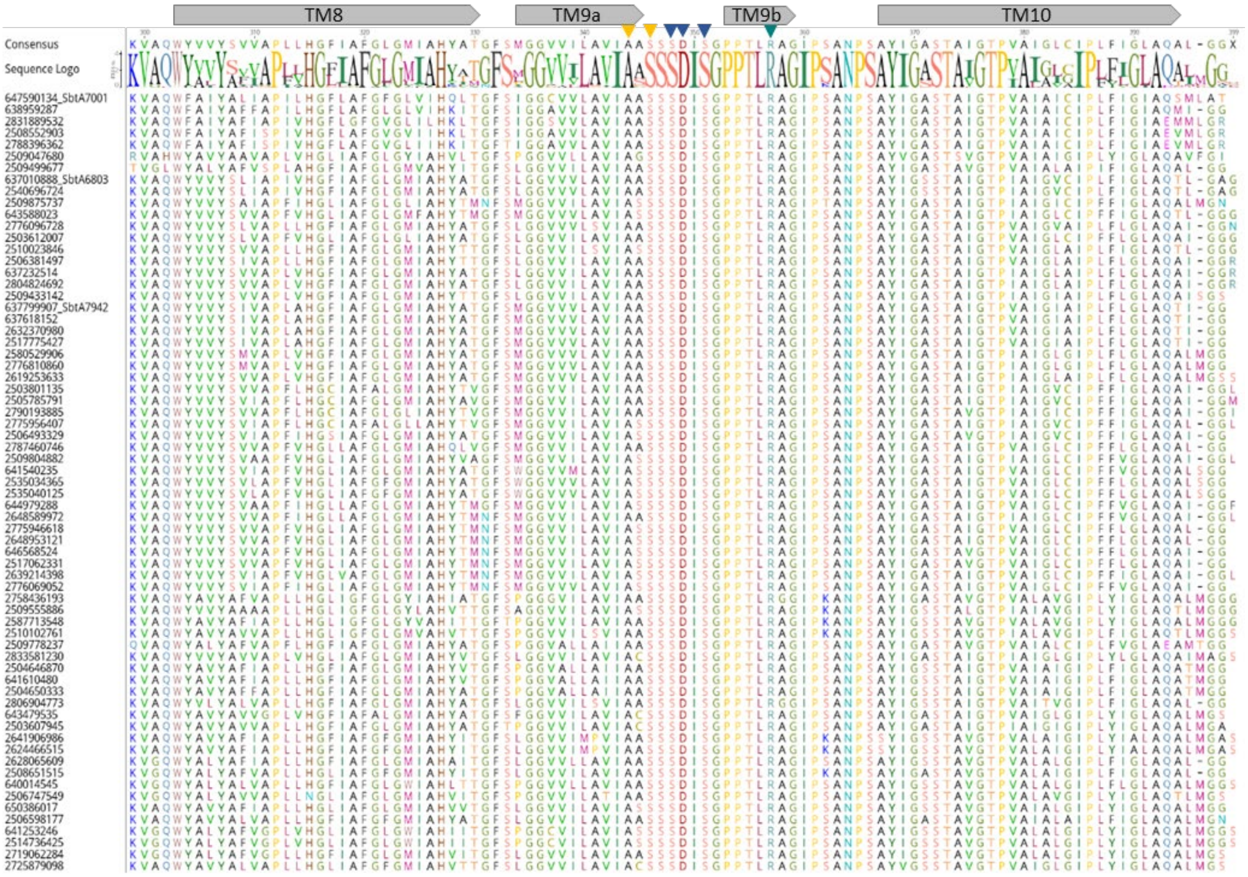

**Fig. S10. Amino acid alignment of SbtA proteins homologs ordered by similarity to $SbtA_{7001}$.**
Multiple pairwise alignments of 67 SbtA proteins homologs (≥ 60% identity). The consensus sequence of the most common residues at each position and the sequence logo are shown above the alignments. Protein identifiers are the gene IDs from IMG (Table S1), and SbtA proteins most relevant to this study are labelled. Predicted transmembrane helices (TM; grey arrows) and residues coordinating the $Na^+$ ion (yellow triangles), $HCO_3^-$ (blue triangles) or implicated in interaction with SbtB (green triangles) are marked as in Fig. S9A.

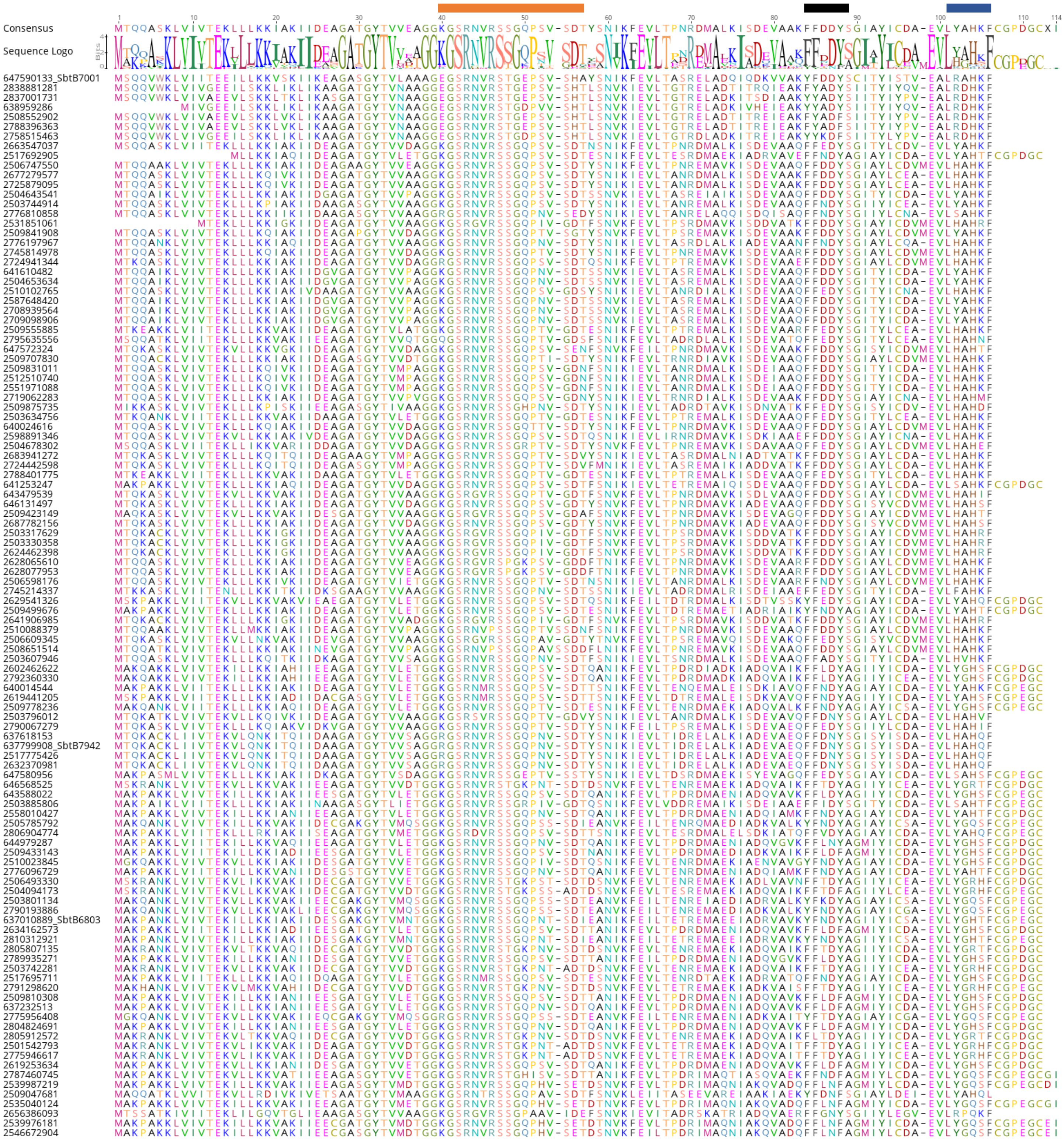


**Fig. S11. Amino acid alignment of SbtB protein homologs ordered by similarity to $SbtB_{7001}$.**
Multiple pairwise sequence alignments of 110 SbtB protein homologs (≥ 50% identity). The consensus sequence comprised of the most common residues at each position and the sequence logo are shown above. Protein identifiers are the gene ID from IMG (Table S2). The SbtA proteins relevant to this study are denoted. The B-, T- and C-loops are indicated by orange, black and blue bars, respectively. Note that the C-terminal extension (CGPxGC) present in $SbtB_{6803}$ which has been suggested to function as redox-sensor is absent from both $SbtB_{7001}$ and $SbtB_{7942}$. This supports the notion that a variety sensory and regulatory mechanisms may have evolved for SbtB proteins in different cyanobacterial species.

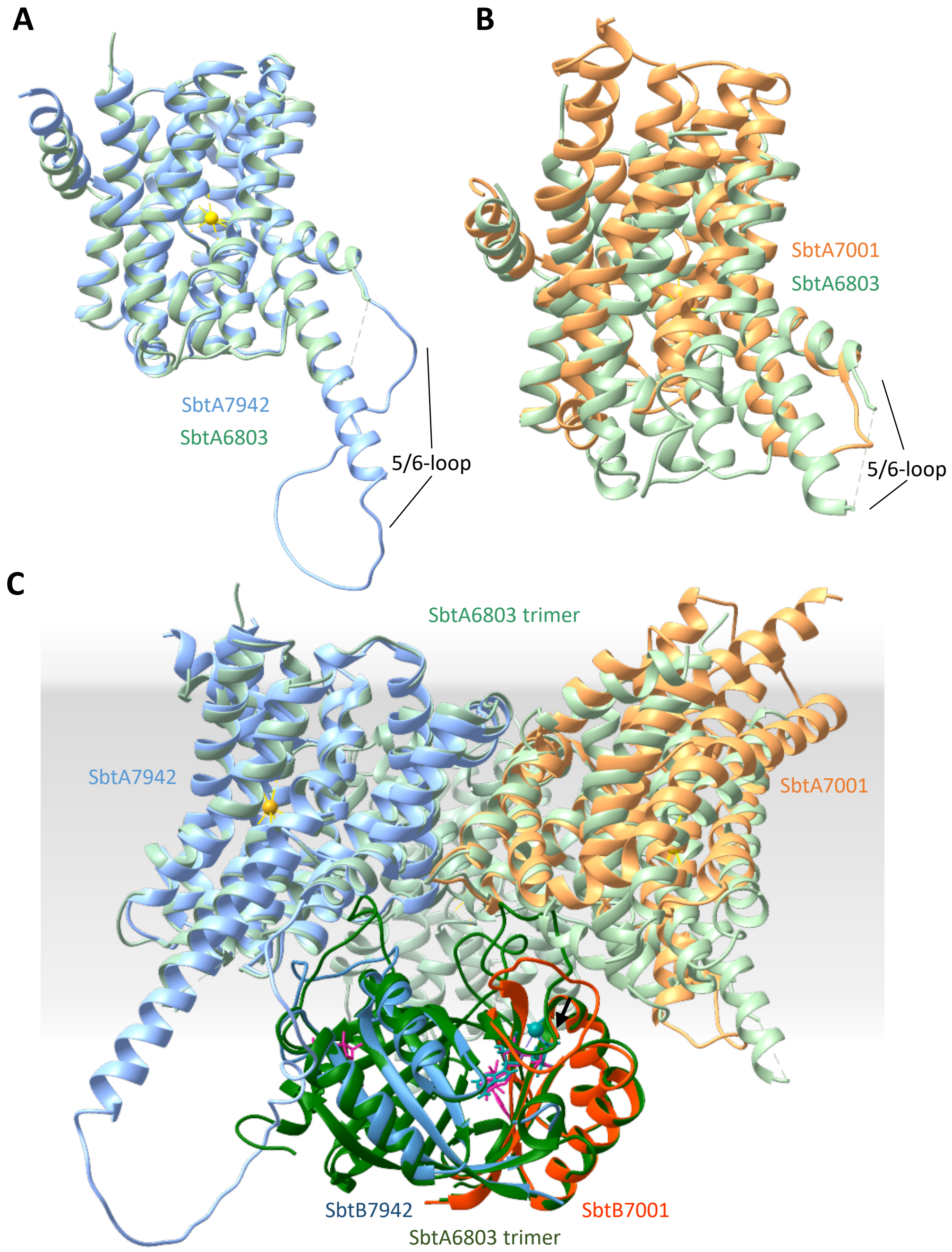


**Fig. S12. Structural comparison of the overall fold of SbtA and SbtB proteins from *Synechococcus elongatus* PCC7942, *Cyanobium sp.* PCC7001 and *Synechocystis sp.* PCC6803.** The three-dimensional homology models of $SbtA_{7942}$ (**A**) and $SbtA_{7001}$ (**B**) were superimposed on the cryo-EM structure of $SbtA_{6803}$ chain C (PDB: 7EGK). Both $SbtA_{7001}$ and $SbtA_{7942}$ show very similar folds to $SbtA_{6803}$. The active site is indicated by the $Na^{+}$ ion (yellow sphere) coordinated with SbtA6803. The variable 5/6 loop was not resolved in the $SbtA_{6803}$, and therefore the 5/6 loops were modelled with lower confidence for $SbtA_{7001}$ and $SbtA_{7942}$. (**C**) The three-dimensional structures of $SbtA_{7001}$, $SbtA_{7942}$, $SbtB_{7001}$ and $SbtB_{7942}$ were superimposed on the crystal structure of the $SbtA_{6803}$ trimer associated with the $SbtB_{6803}$ trimer (PDB: 7EGK) harbouring AMP as ligand (magenta), which illustrates that, based on structural homology, the three SbtA-SbtB pairs may form highly similar trimeric functional units. T-loop conformations of the SbtB proteins are consistent with SbtA-SbtB binding results of this work. The extended (open) conformation of the T-loop of $SbtB_{6803}$ AMP (green) inserts into the gap between the SbtA core and outer scaffold, consistent with biochemical data on SbtA-SbtB binding from *in vitro* assays. The T-loop of $SbtB_{7001}$ $Ca^{2+}$ ATP (dark orange; PDB: 6MM2) assumes a rigid closed position covering the SbtB nucleotide binding cleft which sterically impedes T-loop interaction with SbtA, also consistent with our observations that high levels of $Ca^{2+}$ATP and ATP prevent SbtA:SbtB association.

**Table S1. List of gene identifiers for SbtA protein homologs aligned in Fig. S10.**

| IMG Gene ID[a] | Locus_tag[a] (Protein) | Species |
|---|---|---|
| 647590134 | CPCC7001_1784 (**SbtA$_{7001}$**) | **Cyanobium sp. PCC 7001** |
| 637799907 | Synpcc7942_1475 (**SbtA$_{7942}$**) | **Synechococcus elongatus PCC 7942** |
| 637010888 | slr1512 (**SbtA$_{6803}$**) | **Synechocystis sp. PCC 6803** |
| 637232514 | all2134 | Nostoc sp. PCC 7120 |
| 637618152 | syc2461_c | Synechococcus elongatus PCC 6301 |
| 638959287 | WH5701_01945 | Synechococcus sp. WH5701 |
| 640014545 | L8106_14560 | Lyngbya sp. PCC 8106 |
| 641253246 | AM1_4164 | Acaryochloris marina MBIC11017 |
| 641540235 | MAE_62090 | Microcystis aeruginosa NIES-843 |
| 641610480 | SYNPCC7002_A0470 | Synechococcus sp. PCC 7002 |
| 643479535 | PCC7424_1268 | Gloeothece citriformis PCC 7424 |
| 643588023 | Cyan7425_5063 | Cyanothece sp. PCC 7425 |
| 644979288 | Cyan8802_1280 | Rippkaea orientalis PCC 8802 |
| 646568524 | Ava_3027 | Anabaena variabilis ATCC 29413 |
| 650386017 | NIES39_E03120 | Arthrospira platensis NIES-39 |
| 2503607945 | GEI7407_1945 | Geitlerinema sp. PCC 7407 |
| 2503612007 | Chro_1748 | Chroococcidiopsis thermalis PCC 7203 |
| 2503801135 | Sta7437_3040 | Stanieria cyanosphaera PCC 7437 |
| 2504646870 | SYNPCC7117DRAFT_00005270 | Synechococcus sp. PCC 7117 |
| 2504650333 | SYNPCC8807DRAFT_00005370 | Synechococcus sp. PCC 8807 |
| 2505785791 | Chr6712_2034 | Chroococcidiopsis sp. PCC 6712 |
| 2506381497 | NIVACYA15_00004980 | Planktothrix agardhii NIVA-CYA 15 |
| 2506493329 | Ana7108_3473 | Anabaena sp. PCC 7108 |
| 2506598177 | Spi9445_1079 | Spirulina subsalsa PCC 9445 |
| 2506747549 | Syn7336_2246 | Synechococcus sp. PCC 7336 |
| 2508552903 | Cyagr_2515 | Cyanobium gracile PCC 6307 |
| 2508651515 | Xen7305DRAFT_00036140 | Xenococcus sp. PCC 7305 |
| 2509047680 | ThimaDRAFT_1453 | Thiocapsa marina 5811, DSM 5653 |
| 2509433142 | Mic7113_1307 | Microcoleus sp. PCC 7113 |
| 2509499677 | Pro9006DRAFT_1168 | Prochlorothrix hollandica PCC 9006 |
| 2509555886 | Dacsa_3500 | Dactylococcopsis salina PCC 8305 |
| 2509778237 | Lepto7104DRAFT_5458 | Nodosilinea nodulosa PCC 7104 |
| 2509804882 | LepboDRAFT_4224 | Leptolyngbya boryana PCC 6306 |
| 2509875737 | Syn6308DRAFT_2444 | Geminocystis herdmanii PCC 6308 |
| 2510023846 | Cal7102DRAFT_01206 | Calothrix desertica PCC 7102 |
| 2510102761 | Gei7105DRAFT_4016 | Geitlerinema sp. PCC 7105 |
| 2514736425 | ACCM5_010100012349 | Acaryochloris sp CCMEE 5410 |
| 2517062331 | PCC9339DRAFT_03188 | Fischerella sp. PCC 9339 |
| 2517775427 | synDRAFT_01697 | Synechococcus leopoliensis UTEX 625a |
| 2535034365 | | Microcystis aeruginosa PCC 9808 |
| 2535040125 | | Microcystis aeruginosa PCC 9806 |
| 2540696724 | BEST7613_2847 | Bacillus subtilis BEST7613 |
| 2580529906 | BG1DRAFT_02234 | Synechococcus sp. NKBG15041c |
| 2587713548 | HLSNC16_00948 | Phormidium sp. OSCR GFM |
| 2619253633 | Ga0060318_1481020 | Tolypothrix bouteillei Iicb1 |
| 2624466515 | Ga0064116_101453 | Crocosphaera watsonii WH 8502 |
| 2628065609 | Ga0079929_100413 | Roseofilum sp. BLZ4_bin2 |
| 2632370980 | Ga0069299_132184 | Synechococcus sp. UTEX 2973 |
| 2639214398 | Ga0099329_101977 | Planktothricoides sp. SR001 |
| 2641906986 | Ga0081899_16876 | Crocosphaera watsonii WH 8502 |
| 2648589972 | Ga0077703_1084379 | Scytonema millei VB511283 |
| 2648953121 | Ga0072975_102525 | Tolypothrix sp. PCC 7601 |
| 2719062284 | Ga0174825_114265 | Moorea producens PAL-8-15-08-1 |
| 2725879098 | Ga0126230_16858 | Anabaena sp. ATCC 33047 |
| 2758436193 | Ga0225988_11931 | Halomicronema hongdechloris C2206 |
| 2775946618 | Ga0263755_110933 | Microchaete diplosiphon NIES-3275 |
| 2775956407 | Ga0263656_172662 | Chondrocystis sp. NIES-4102 |

| | | |
|---|---|---|
| 2776069052 | Ga0263379_123731 | Sphaerospermopsis kisseleviana NIES-73 |
| 2776096728 | Ga0262663_111374 | Nodularia sp. NIES-3585 |
| 2776810860 | Ga0264152_141819 | Synechococcus sp. NIES-970 |
| 2787460746 | Ga0302962_10599 | Elainella saxicola E1 |
| 2788396362 | Ga0303538_101391 | Aphanothece cf. minutissima CCALA 015 |
| 2790193885 | Ga0303530_103124 | Pleurocapsa sp. CCALA 161 |
| 2804824692 | Ga0263997_15134 | Trichormus variabilis NIES-23 |
| 2806904773 | Ga0316382_139810 | Rubidibacter sp. SB-MAG 54 |
| 2831889532 | Ga0336076_95 | Vulcanococcus limneticus LL |
| 2833581230 | Ga0335515_2769 | Thermosynechococcus elongatus PKUAC-SCTE542 |

[a] Gene ID and Locus tags in accordance with IMG. SbtA proteins most relevant to this study are in bold.

**Table S2. List of gene identifiers for SbtB protein homologs aligned in Fig. S11.**

| IMG Gene ID[a] | Locus_tag [a] (Protein) | Species |
|---|---|---|
| 647590133 | CPCC7001_1671 (**$SbtB_{7001}$**) | **Cyanobium sp. PCC 7001** |
| 637799908 | Synpcc7942_1476 (**$SbtB_{7942}$**) | **Synechococcus elongatus PCC 7942** |
| 637010889 | slr1513 (**$SbtB_{6803}$**) | **Synechocystis sp. PCC 6803** |
| 637232513 | all2133 | Nostoc sp. PCC 7120 |
| 637618153 | syc2462_d | Synechococcus elongatus PCC 6301 |
| 638959286 | WH5701_01940 | Synechococcus sp. WH5701 |
| 640014544 | L8106_14555 | Lyngbya sp. PCC 8106 |
| 640024616 | N9414_20315 | Nodularia spumigena CCY9414 |
| 641253247 | AM1_4165 | Acaryochloris marina MBIC11017 |
| 641610482 | SYNPCC7002_A047 | Synechococcus sp. PCC 7002 |
| 643479539 | PCC7424_1272 | (Gloeothece citriformis) Cyanothece sp. PCC 7424 |
| 643588022 | Cyan7425_5062 | Cyanothece sp. PCC 7425 |
| 644979287 | Cyan8802_1279 | (Rippkaea orientalis) Cyanothece sp. PCC 8802 |
| 646131497 | AplaP_010100019251 | Arthrospira platensis Paraca |
| 646568525 | Ava_3028 | Anabaena variabilis ATCC 29413 |
| 647572324 | MC7420_8214 | Coleofasciculus chthonoplastes PCC 7420 |
| 647580956 | S7335_623 | Synechococcus sp. PCC 7335 |
| 2501542793 | fdiDRAFT38940 | Tolypothrix sp. PCC 7601 |
| 2503317629 | CWat_WH0401_DRAFT_00006160 | Crocosphaera watsonii WH 0401 |
| 2503330358 | CWat_WH8501_draft2_00012570 | Crocosphaera watsonii WH 8501 |
| 2503607946 | GEI7407_1946 | Geitlerinema sp. PCC 7407 |
| 2503634756 | PCC7418_0057 | Halothece sp. PCC 7418 |
| 2503742281 | Nos7107_3598 | Nostoc sp. PCC 7107 |
| 2503744914 | Cyan10605_0668 | Cyanobacterium aponinum PCC 10605 |
| 2503796012 | Glo7428_3433 | Gloeocapsa sp. PCC 7428 |
| 2503801134 | Sta7437_3039 | Stanieria cyanosphaera PCC 7437 |
| 2503885806 | Lepto7376_0211 | Leptolyngbya sp. PCC 7376 |
| 2504094173 | Cal6303_1189 | Calothrix parietina PCC 6303 |
| 2504643541 | SYNPCC7003DRAFT_00005420 | Synechococcus sp. PCC 7003 |
| 2504653634 | SYNPCC73109DRAFT_00005450 | Synechococcus sp. PCC 73109 |
| 2504678302 | Pse7367_3849 | Pseudanabaena sp. PCC 7367 |
| 2505785792 | Chr6712_2035 | Chroococcidiopsis sp. PCC 6712 |
| 2506493330 | Ana7108_3474 | Anabaena sp. PCC 7108 |
| 2506598176 | Spi9445_1078 | Spirulina subsalsa PCC 9445 |
| 2506609345 | Spi6313_1704 | Spirulina major PCC 6313 |
| 2506747550 | Syn7336_2247 | Synechococcus sp. PCC 7336 |
| 2508552902 | Cyagr_2514 | Cyanobium gracile PCC 6307 |
| 2508651514 | Xen7305DRAFT_00036130 | Xenococcus sp. PCC 7305 |
| 2509047681 | ThimaDRAFT_1454 | Thiocapsa marina 5811, DSM 5653 |
| 2509423149 | Oscil6304_3942 | Oscillatoria acuminata PCC 6304 |
| 2509433143 | Mic7113_1308 | Microcoleus sp. PCC 7113 |
| 2509499676 | Pro9006DRAFT_1167 | Prochlorothrix hollandica PCC 9006 |
| 2509555885 | Dacsa_3499 | Dactylococcopsis salina PCC 8305 |
| 2509707830 | Pleur7313DRAFT_01818 | Pleurocapsa sp. PCC 7319 |
| 2509778236 | Lepto7104DRAFT_5457 | Nodosilinea nodulosa PCC 7104 |
| 2509810308 | Nos7524_2739 | Nostoc sp. PCC 7524 |
| 2509831011 | CfriDRAFT_00028170 | Chlorogloeopsis fritschii PCC 6912 |
| 2509841908 | Lepto7375DRAFT_1925 | Leptolyngbya sp. PCC 7375 |
| 2509875735 | Syn6308DRAFT_2442 | (Geminocystis herdmanii) Synechocystis sp. PCC 6308 |
| 2510023845 | Cal7102DRAFT_01205 | Calothrix desertica PCC 7102 |
| 2510088379 | Riv7116_3233 | Rivularia sp. PCC 7116 |
| 2510102765 | Gei7105DRAFT_4020 | Geitlerinema sp. PCC 7105 |
| 2512510740 | Cfri_03095 | Chlorogloeopsis fritschii PCC 6912 |
| 2517692905 | LEP6406DRAFT_2363 | Leptolyngbya sp. PCC 6406 |
| 2517695711 | LEP6406DRAFT_5170 | Leptolyngbya sp. PCC 6406 |
| 2517775426 | synDRAFT_01696 | Synechococcus leopoliensis UTEX 625a |
| 2531851061 | CWATWH0003_5162 | Crocosphaera watsonii WH 0003 |
| 2535040124 | | Microcystis aeruginosa PCC 9806 |

| | | |
|---|---|---|
| 2539976181 | | Microcystis aeruginosa PCC 7941 |
| 2539987219 | | Microcystis aeruginosa PCC 9443 |
| 2546672904 | MAESPC_02398 | Microcystis aeruginosa SPC777 |
| 2551971088 | UYCDRAFT_06647 | Chlorogloeopsis fritschii PCC 6912 |
| 2558010427 | Osccy1DRAFT_03495 | Leptolyngbya sp. JSC-1 |
| 2587648420 | BG2DRAFT_01749 | Synechococcus sp. NKBG 042902 |
| 2598891346 | Ga0059189_02159 | Myxosarcina sp. GI1 |
| 2602462622 | Ga0039500_101179 | Planktothrix agardhii NIVA-CYA 15 |
| 2619253634 | Ga0060318_1481021 | Tolypothrix bouteillei Iicb1 |
| 2619441205 | Ga0062155_104430 | Leptolyngbya sp. KIOST-1 |
| 2624462398 | Ga0064117_12042 | Crocosphaera watsonii WH 8501 |
| 2628065610 | Ga0079929_100414 | Roseofilum sp. BLZ4_bin2 |
| 2628077953 | Ga0079931_15633 | Roseofilum sp. LKpool_bin4 |
| 2629541326 | Ga0078341_12711765 | Aphanocapsa montana BDHKU210001 |
| 2632370981 | Ga0069299_132185 | Synechococcus sp. UTEX 2973 |
| 2634162573 | Ga0080671_6209 | Nostoc sp. 996 |
| 2641906985 | Ga0081899_16875 | Crocosphaera watsonii WH 8502 |
| 2656386093 | Ga0111153_11066 | Gammaproteobacteria bacterium SG8_31 |
| 2663547037 | Ga0115370_115168 | Geitlerinema sp. PCC 9228 re-assembly |
| 2677279577 | Ga0126231_16554 | Anabaena sp. 4-3 |
| 2683941272 | Ga0123818_111979 | Leptolyngbya sp. O-77 |
| 2687782156 | Ga0133377_111972 | Arthrospira platensis YZ |
| 2708939564 | Ga0154051_0543 | Synechococcus sp. 7002 |
| 2709098906 | Ga0154052_1261 | Synechococcus sp. OG1 |
| 2719062283 | Ga0174825_114264 | Moorea producens PAL 15AUG08-1 |
| 2724442598 | Ga0182245_108113 | Oscillatoria sp. 08 |
| 2724941344 | Ga0125915_110759 | Mastigocoleus testarum BC008 |
| 2725879095 | Ga0126230_16855 | Anabaena sp. ATCC 33047 |
| 2745214337 | Ga0132956_102022 | Phormidium willei BDU 130791 |
| 2745814978 | Ga0133135_10867 | Nodularia spumigena CENA596 |
| 2758515463 | Ga0226272_111506 | Cyanobium sp. NIES-981 |
| 2775946617 | Ga0263755_110932 | Microchaete diplosiphon NIES-3275 |
| 2775956408 | Ga0263656_172663 | Chondrocystis sp. NIES-4102 |
| 2776096729 | Ga0262663_111375 | Nodularia sp. NIES-3585 |
| 2776197967 | Ga0263577_114559 | Calothrix sp. NIES-3974 |
| 2776810858 | Ga0264152_141817 | Synechococcus sp. NIES-970 |
| 2787460745 | Ga0302962_10598 | Elainella saxicola E1 |
| 2788396363 | Ga0303538_101392 | Aphanothece cf. minutissima CCALA 015 |
| 2788401775 | Ga0303139_106612 | Euhalothece sp. PCC 7418 |
| 2789935271 | Ga0303534_108719 | Cyanosarcina cf. burmensis CCALA 770 |
| 2790067279 | Ga0272292_112316 | Cyanobacterium sp. HL-69 |
| 2790193886 | Ga0303530_103125 | Pleurocapsa sp. CCALA 161 |
| 2791298620 | Ga0272299_132323 | Nostoc sp. CENA543 |
| 2792360330 | Ga0308451_117115 | Planktothrix paucivesiculata PCC 9631 |
| 2795635556 | Ga0310153_10333 | Euhalothece sp. SG1_44_8 |
| 2804824691 | Ga0263997_15133 | Trichormus variabilis NIES-23 |
| 2805807135 | Ga0309075_10103 | Fischerella thermalis CCMEE 5318 |
| 2805912572 | Ga0309080_14417 | Fischerella thermalis CCMEE 5205 |
| 2806904774 | Ga0316382_139811 | Rubidibacter sp. SB-MAG 54 |
| 2810312921 | Ga0325128_122786 | Synechocystis sp. PCC 6714 |
| 2837001731 | Ga0339340_2154 | Synechococcus sp. MW101C3 |
| 2838881281 | Ga0339341_443 | Synechococcus sp. BO 8801 |

[a] Gene ID and Locus tags in accordance with IMG. SbtB proteins most relevant to this study are in bold.
